# Detecting and phasing minor single-nucleotide variants from long-read sequencing data

**DOI:** 10.1101/2020.09.25.314252

**Authors:** Zhixing Feng, Jose Clemente, Brandon Wong, Eric E. Schadt

## Abstract

Cellular genetic heterogeneity is common in many biological conditions including cancer, microbiome, co-infection of multiple pathogens. Detecting and phasing minor variants, which is to determine whether multiple variants are from the same haplotype, play an instrumental role in deciphering cellular genetic heterogeneity, but are still difficult because of technological limitations. Recently, long-read sequencing technologies, including those by Pacific Biosciences and Oxford Nanopore, have provided an unprecedented opportunity to tackle these challenges. However, high error rates make it difficult to take full advantage of these technologies. To fill this gap, we introduce iGDA, an open-source tool that can accurately detect and phase minor single-nucleotide variants (SNVs), whose frequencies are as low as 0.2%, from raw long-read sequencing data. We also demonstrated that iGDA can accurately reconstruct haplotypes in closely-related strains of the same species (divergence *≥* 0.011%) from long-read metagenomic data. Our approach, therefore, presents a significant advance towards the complete deciphering of cellular genetic heterogeneity.

## Introduction

Cellular genetic heterogeneity is prevalent in multiple biological conditions. For example, the microbiome contains multiple bacterial species with distinct genomes, and patients with infections may carry multiple bacterial strains. Likewise, in cancer, tumors are typically characterized by multiple cell types and cell lineages with different genomes. Deconvoluting such complex cellular genetic heterogeneity is critical to basic biology and precision medicine. Minor variants, which are defined as the variants with frequencies lower than 10% in a cell population, play a central role in deciphering cellular genetic heterogeneity. Short-read genome sequencing can effectively characterize a large number of cells simultaneously but cannot phase minor variants directly due to the limitation of read length, which is generally under 300 bp^1^. Long-read sequencing, on the other hand, can be used to overcome this limitation. The latest long-read sequencing technologies, including those by Pacific Biosciences (PacBio) and Oxford Nanopore (ONT), enable sequencing more than 100 billion bases in a single run and yield reads with lengths that can exceed 10 kb^2–4^. These advantages make it feasible to adopt long-read sequencing to study cellular genetic heterogeneity in the microbiome, bacterial co-infection, and cancer in finer details. Because of its long read-length and high throughput, long-read sequencing has the potential to be applied to detect and phase minor variants at the single-molecule level without amplification. However, the error rate of raw long-read sequencing data is usually higher than 10%^1,3^, and makes it difficult to detect variants whose frequency is lower than the sequencing error rate.

Most of the existing methods to detect minor SNVs are based on short-read sequencing data^5–14^. The vast majority of these methods scan the reference genome and detect SNVs or other variants locus-by-locus. These methods cannot be used for long-read sequencing data because they are based on the error pattern of short-read sequencing data, which is different from long-read sequencing data. Researchers have also tried to leverage the information of multiple SNVs to increase detection accuracy. V-Phaser and V-Phaser2^15,16^, which were designed for short-read sequencing data, use the joint probability of SNV pairs to detect SNVs. However, to avoid combinatorial explosion, they only use the joint probability of two SNVs. We will discuss the limitations of such a restriction for long-read sequencing and demonstrate how it leads to false negatives in Results.

There are several methods designed specifically to detect variants from long-read sequencing data. The GenomicConsensus module (https://github.com/PacificBiosciences/GenomicConsensus) developed by PacBio generates a consensus sequence from the aligned PacBio reads and compares it to the reference genome to identify variants. Nanopolish^17^ is a variant caller designed specifically for ONT data, and Clairvoyante^18^ is a deep-learning based tool for Illumina, PacBio, and ONT data. These methods assume that samples only have one or two haplotypes and therefore cannot be applied to detect minor variants. MinorSeq (https://github.com/PacificBiosciences/minorseq), developed by PacBio, is designed to detect minor variants but requires its input to be Circular Consensus Sequencing (CCS) reads^19^. CCS is a special protocol of PacBio sequencing, which sequences each DNA molecule multiple times to increase accuracy. However, CCS reduces read length by 10 to 20 fold to achieve low error rates, and read length is critical to phasing minor SNVs. Recently, several tools have been developed to detect variants by leveraging haplotype information from long-read sequencing data^20–22^, but they also assume that the number of haplotypes is one or two. Thus, they cannot be applied to detect minor variants. To our best knowledge, there is currently no tool available to detect minor SNVs from raw data of long-read sequencing.

There are several short-read based methods available to phase minor SNVs^23–29^. These methods cluster the reads locally and phase distant SNVs, whose distances are longer than read length, using statistical models with strong assumptions. The major limitation of these methods is that they phase distant minor SNVs only based on indirect evidence because the read length is too short to span over the distant SNVs. This limitation can be overcome by using long-read sequencing data. The existing haplotyping methods for long-read sequencing data^20–22^ assume there are only one or two haplotypes, and thus cannot be used to phase minor SNVs because the number of haplotypes is unknown.

To address the challenges of detecting and phasing minor SNVs, we developed a novel tool named iGDA (*in vivo* Genome Diversity Analyzer), which can accurately detect and phase minor SNVs, whose frequencies are as low as 0.2%. To detect minor SNVs, iGDA leverages the information of multiple loci without restricting the number of dependent loci, and uses a novel algorithm, Random Subspace Maximization (RSM), to overcome the issue of combinatorial explosion. To phase minor SNVs, iGDA uses a novel algorithm, Adaptive Nearest Neighbor clustering (ANN), which makes no assumption about number of haplotypes. To evaluate the performance of iGDA, we tested it on four pooled longread sequencing datasets. The number of samples pooled in each dataset ranges from 65 to 755. The results demonstrate that iGDA can detect 85.8% to 96.7% of the real SNVs in these datasets at false discovery rate (FDR) lower than 1%. Finally, iGDA can phase minor SNVs at average accuracies range from 90.7% to 98.7%. We also tested iGDA on a pooled long-read metagenomic dataset consisting of 11 *Borrelia burgdorferi* strains and 744 other bacterial species, and discovered that the accuracy of iGDA is sufficient to reconstruct haplotypes in closely-related conspecific strains (strains belonging to the same species) only using one reference genome. The divergences between the distinguishable conspecific strains are as low as 0.011%. These results shed light on tackling a number of challenges such as extracting strain-resolved genome sequences from long-read metageonmic data and identifying multiple strains in co-infection.

## Results

### Detecting minor SNVs by leveraging information of multiple loci

The major challenge of detecting minor SNVs is to distinguish between real SNVs and sequencing errors. It is especially difficult for raw data of long-read sequencing technologies, including those by PacBio and ONT, because they have relatively high error rates. However, we could leverage the fact that long reads can cover multiple SNVs to substantially increase detection accuracy. Intuitively, assuming that sequencing errors are independent, multiple sequencing errors are unlikely to repeatedly occur together on the same read. For example, in a pooled PacBio sequencing dataset consisting of 186 *Bordetella* spp. samples (Figure 1A), the substitutions from the five marked loci occur together on 28 reads and there are 23,432 reads covering these five loci. The observed joint probability that these five substitutions occur together on the same read is 28*/*23432 = 0.00119, while the expected joint probability is less than 0.1^5^ = 0.00001 because substitution error rate of raw PacBio reads is less than 0.1 on this dataset (Figure 1B). The observed joint probability is over 100 times higher than the expected joint probability, so it is very likely that some of the five substitutions are real SNVs. However, the substitution rates of these five SNVs are 0.00569, 0.00845, 0.00748, 0.00960, and 0.00915 respectively and it is difficult to distinguish them from sequencing errors only based on substitution rate (Figure 1B). Based on these observations, we propose a novel framework that uses conditional substitution rate instead of substitution rate to detect SNVs. In this framework, for each substitution, we adopt the maximal probability of observing the substitution conditional on observing substitutions at *p* other loci, defined as maximal conditional substitution rate, to detect whether the substitution is a real SNV. We call these *p* loci dependent loci. However, as the *p* dependent loci are unknown, it is infeasible to enumerate all combinations of these *p* loci to calculate the maximal conditional substitution rate due to high computational cost. As *p* is unknown, the number of combinations is about 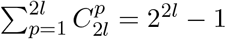 for each locus if the average read length is *l*. We propose a novel algorithm called Random Subspace Maximization (RSM) to estimate the maximal conditional substitution rate efficiently (Figure 2A-C) (details are in Methods). As shown in Figure 1C, on the *Bordetella* spp. data, the real SNVs and the sequencing errors are highly distinguishable based on the maximal conditional substitution rate calculated by the RSM algorithm.

**Figure 1:**
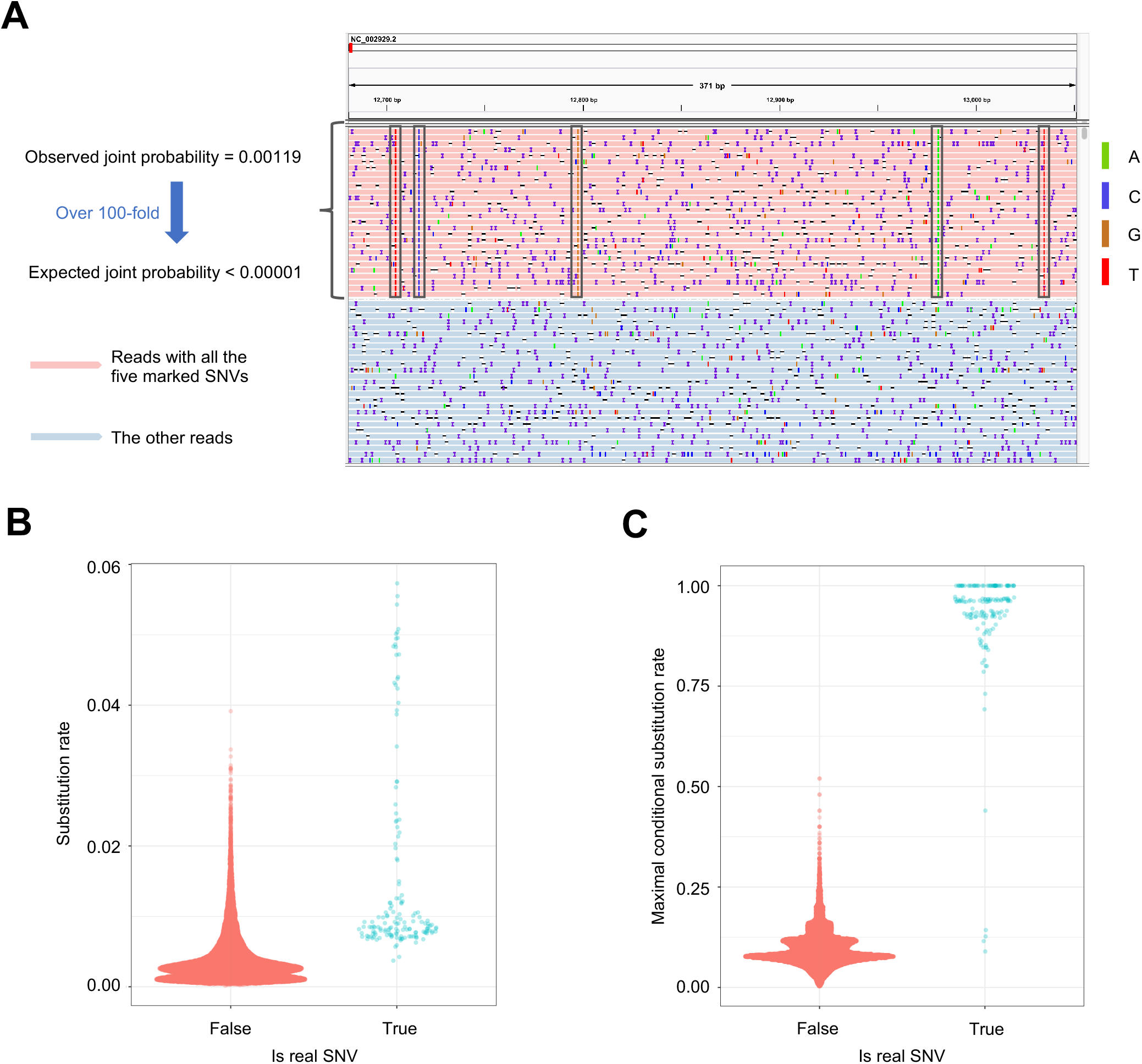
SNVs are dependent on each other. **A**, An IGV (Integrative Genomics Viewer)^30^ snapshot demonstrating how to use the information of multiple loci to increase detection accuracy of SNVs. The number of reads containing the five SNVs marked by black boxes is 28 and the number of reads covering the five SNVs is 23,432. The observed and expected joint probabilities of the five SNVs are shown to the left of the IGV snapshot. Some reads are not shown in the figure due to the limit of figure size. **B**, The distribution of substitution rate on the *Bordetella* spp. data. No outlier is removed in the Sina plot. **C**, The distribution of maximal conditional substitution rate estimated by the RSM algorithm on the *Bordetella* spp. data. No outlier is removed in the Sina plot.

**Figure 2:**
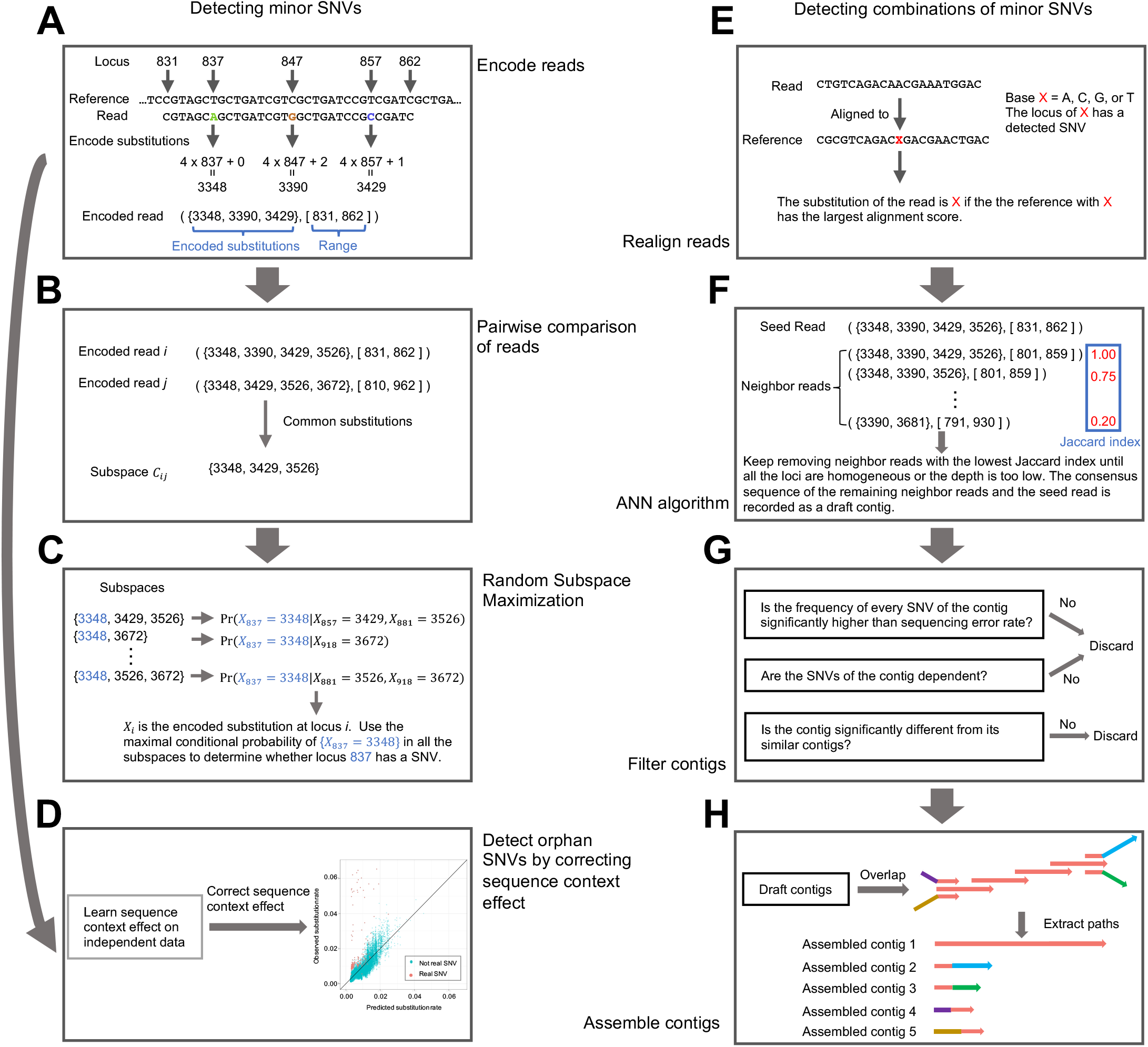
The main steps of iGDA. Details are in the Methods section

It is very important to note that the number of dependent loci *p* should not be fixed. Figure S1 shows an example that fixing *p* can induce false negatives. In this example, the substitution at the locus 1 is independent with the substitutions at locus 2 and locus 3 respectively, but highly dependent on the combination of the substitutions at locus 2 and locus 3. Thus, the SNV at locus 1 is difficult to be detected if *p* is fixed to 1, but is easy to be detected if there is no restriction on *p*. The existing algorithms V-Phaser and V-phaser2^15,16^ were designed to identify minor variants from short-read sequencing data and only leveraged dependence between substitutions at two loci to avoid combinatorial explosion. This is equivalent to fixing *p* to 1, and making these algorithms unable to detect the SNVs in Figure S1. The proposed RSM algorithm has no restriction on *p* and can avoid combinatorial explosion.

If a SNV is the only SNV in the genome, we call it an orphan SNV. The proposed framework that uses conditional substitution rate to detect SNVs cannot detect orphan SNVs because its basic assumption is that there are multiple real SNVs in the same genome. We propose a single-locus based algorithm to overcome this limitation (Figure 2D). We discovered that substitution error rate is very different from locus to locus and it is highly predictable by sequence context (Figure 3). We trained a gradient boosting model^31^ on independent public data and predicted substitution error rate for each locus. We then adopted a likelihood ratio test to compare the observed substitution rate to the predicted substitution error rate and reported a SNV if they are significantly different (details are in Methods).

**Figure 3:**
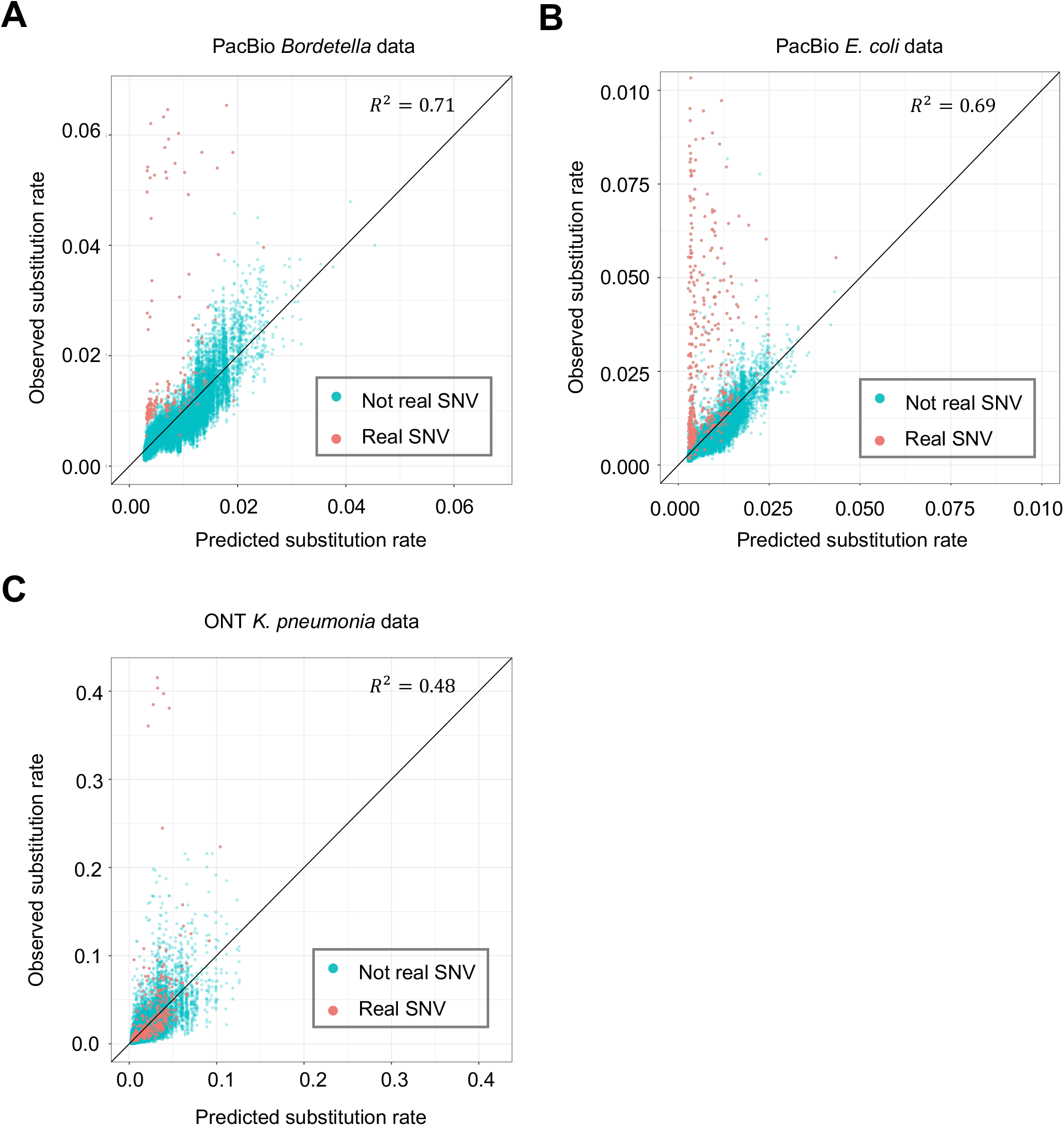
Predicting substitution error rate by sequence-context-effect model trained on independent data. **A**, Prediction of substitution error rate on the PacBio *Bordetella* spp. data. **B**, Prediction of substitution error rate on the PacBio *E. coli* data. **C**, Prediction of substitution error rate on the ONT *k. pneumoniae* data with DNA methylation masked.

### Phasing minor SNVs

Intuitively, the reads of the same genome should be clustered together and the consensus sequence of each cluster can be used to phase minor SNVs. Herein, we propose a novel algorithm called Adaptive Nearest Neighbor (ANN) to cluster the reads and the consensus sequence of each cluster is called a draft contig (Figure 2E and Figure 2F) (details are in Methods). To reduce noise, loci with no detected SNVs are masked before applying ANN algorithm. A major advantage of ANN algorithm is that it can estimate the number of clusters automatically while clustering the reads. To reduce false positive rate of the draft contigs, we adopted a two-step filter to remove unreliable draft contigs (Figure 2G). Intuitively, the SNVs in the same draft contig should be dependent with each other and the difference between two similar draft contigs should be statistically significant.

The lengths of the draft contigs are usually smaller than genome size. To maximize the range where the minor SNVs can be phased, we assemble the draft contigs using an algorithm inspired by overlap graph^32^ (Figure 2H) (details are in Methods). The assembled draft contigs are called contigs.

### Evaluating performance on pooled PacBio sequencing data

We constructed two datasets to test the accuracy of iGDA. The first dataset is a mixture of PacBio sequencing data of 186 *Bordetella* spp. samples, and the second dataset is a mixture of 155 *Escherichia coli* samples. The datasets have been previously published and their accession IDs in the SRA database (https://www.ncbi.nlm.nih.gov/sra) are listed in Table S1. The average sequencing depths of pooled data are 29,208x for *Bordetella* spp. and 19,175x for *Escherichia coli*. We downloaded the raw data in HDF format from SRA, and filtered the reads by requiring the estimated read quality (rq) greater than 0.75. The estimated read quality were extracted from the native HDF file. Bases with quality value (QV) less than a threshold were masked. We tested four thresholds, 0, 8, 10, and 12, respectively. We aligned the filtered reads to the reference genomes of *Bordetella pertussis* Tohama I (NCBI Reference Sequence ID is NC_002929.2) for the *Bordetella* spp. data and *Escherichia coli* K12 MG1655 (NCBI reference sequence ID is NC_000913.3) for the *Escherichia coli* data by minimap2 (version 2.12)^33^ respectively. To minimize the alignment ambiguity caused by the aligner, we realigned the reads mapped to the negative strand by aligning their reverse complementary sequences. We only retained the reads aligned to the concatenated *rpoB* and *rpoC* region, which is highly conserved. The 1-based coordinates of the reference genomes is [11662, 20018] for *Bordetella pertussis* Tohama I and [4181245, 4189573] for *Escherichia coli* K12 MG1655. We pooled the realigned reads aligned to the concatenated *rpoB* and *rpoC* region for *Bordetella* spp. and *Escherichia coli* respectively to construct the two datasets. To evaluate accuracy of iGDA, we ran PacBio’s genome consensus module (https://github.com/pacificbiosciences/genomicconsensus) on the aligned reads of each sample with default parameters to obtain the consensus genome sequences and SNVs. The union of the SNVs were used as benchmark to evaluate the accuracy of detecting SNVs. The genome sequence of an individual sample is defined as a real contig, and was used to evaluate the accuracy of contigs reported by iGDA. We merged samples (real contigs) with identical SNV profiles and calculated the relative abundances of the merged samples by the ratio between number of reads aligned to each sample and the total number of aligned reads. The relative abundances of the samples distinct from the reference genome range from 0.25% to 3.05% for the *Bordetella* spp. data, and range from 0.30% to 1.92% for the *Escherichia coli* data. The average relative abundances are 0.82% and 0.74% for the *Bordetella* spp. data and the *Escherichia coli* data respectively.

For detecting minor SNVs, we tested three algorithms—a single-locus method (SL), which simply uses substitution rate of each locus to detect SNVs; a context-aware single-locus method (SLC), which uses substitution rate of each locus with correcting sequence-context effect (details are in Methods); and the proposed RSM algorithm—on these two test datasets. The results indicate that RSM algorithm greatly outperforms the two single-locus methods, and achieves a high accuracy (Figure 4A and Figure 4B). With masking bases with QV lower than 8, iGDA detected 96.7% and 85.8% of the real SNVs at false discovery rate (FDR) lower than 1% for the *Bordetella* spp. data and *Escherichia coli* data respectively. Besides, correcting sequence-context effect substantially increases detection accuracy of the single-locus methods. The threshold of base QV also has minor impact on the accuracy. A non-zero threshold increases the accuracy on the *Bordetella* spp. data (Figure 4A), but decreases the accuracy on the *Escherichia coli* data (Figure 4B). This might be because masking bases with low QV removes some sequencing errors but reduces effective sequencing depth.

**Figure 4:**
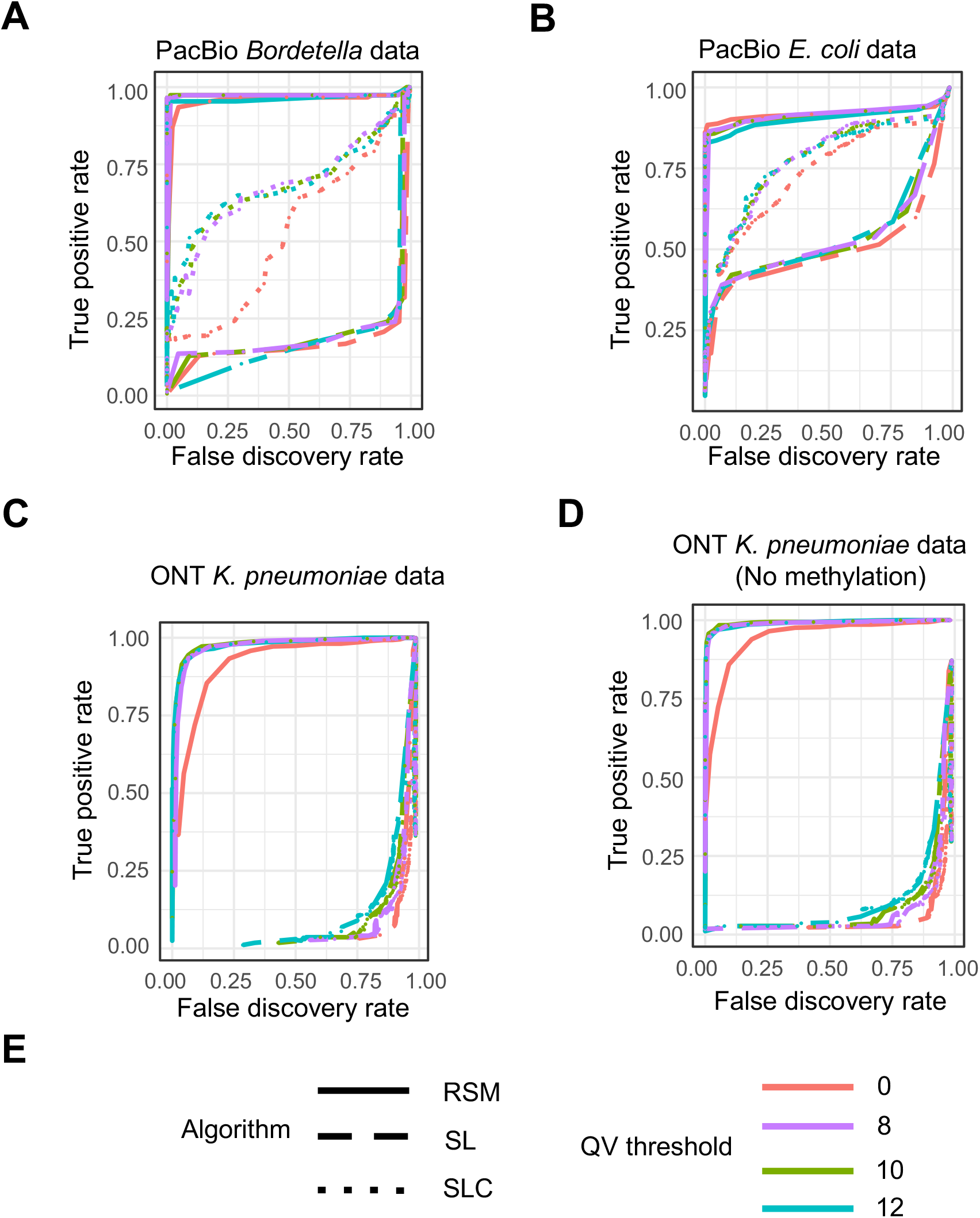
The accuracy of detecting minor SNVs on pooled sequencing data. **A**, The accuracy on PacBio *Bordetella* spp. data. **B**, The accuracy on PacBio *E. coli* data. **C**, The accuracy on ONT *K. pneumoniae* data. **D**, The accuracy on ONT *K. pneumoniae* data with DNA methylation masked. **E**, The legend of subfigures **A**-**D**. RSM = Random Subspace Maximization algorithm, SL = Single-Locus algorithm, SLC = Single-Locus algorithm with correcting sequence-context effect, and QV = Quality Value. True positive rate = number of correctly detected SNVs / number of real SNVs. False discovery rate = 1 - number of correctly detected SNVs / number of detected SNVs.

For phasing minor SNVs, we evaluated the ANN algorithm on these two datasets, where the bases with QV less than 8 were masked. The average accuracies (the maximal Jaccard index^34^ with the real contigs) of the assembled contigs are 98.9% and 98.1% for the *Bordetella* spp. data and *Escherichia coli* data respectively (Figure 5A). Jaccard index between an iGDA-inferred contig and a real contig is the ratio between the number of shared SNVs and the total number of unique SNVs in their overlapped region. The IGV (Integrative Genomics Viewer)^30^ snapshot of the contigs obtained from the *Bordetella* spp. data and the *Escherichia coli* data are shown in Figure 5B and Figure S2. The results show that the iGDA-inferred contigs match the real contigs very well, even for the real contigs with frequencies lower than 1%. In Figure 5B, there are five real contigs that are not detected by our algorithm. One of them has no SNV (the reference genome); two of them only have a single orphan SNV with very low frequency, which is hard for the RSM algorithm to detect; and two of them are highly similar to another genome. The results indicate that the minor SNVs can be phased effectively except for the genomes that have an orphan SNV or are highly similar to another genome.

**Figure 5:**
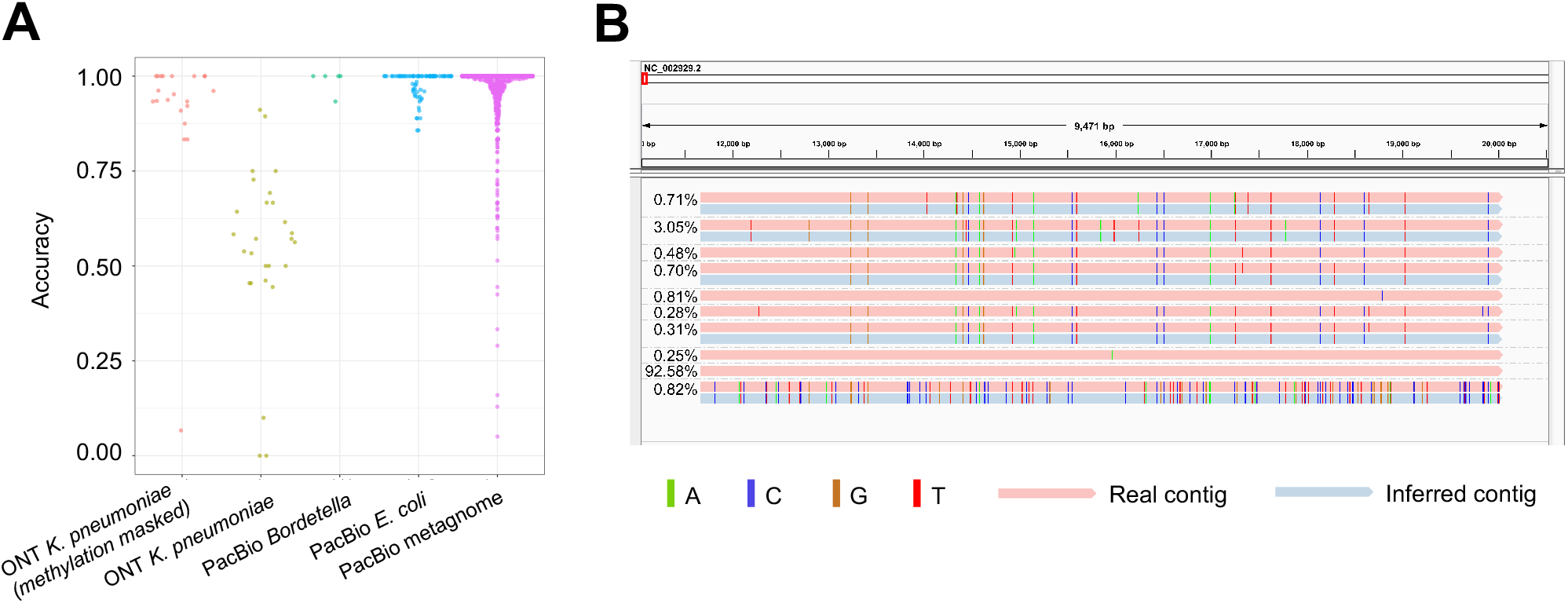
The accuracy of phasing minor SNVs. **A**, The sina plot of accuracy of phasing minor SNVs on the four testing datasets. **B**, The IGV snapshot of the contigs inferred by iGDA on the PacBio *Bordetella* spp. data. An inferred contig is grouped with its most similar real contig (measured by Jaccard index). Relative abundance is shown to the left of each contig.

### Evaluating performance on pooled ONT sequencing data

We tested iGDA on a dataset consisting of a mixture of ONT sequencing data of 65 *Klebsiella pneumoniae* samples. The SRA IDs are listed in Table S2. We downloaded the raw data in fastq format from the SRA database (https://www.ncbi.nlm.nih.gov/sra), filtered and trimmed the reads using fastp^35^. The reads with average quality value (QV) less than 8 were discarded, and the first 50 bp and the last 200 bp were trimmed for each read. Similar to the PacBio data, we used four thresholds, 0, 8, 10, and 12 respectively, to mask bases with low QV. The reads were then aligned to the reference genome of *Klebsiella pneumoniae* subsp. pneumoniae HS11286 (NCBI reference sequence ID is NC_016845.1). We realigned the reads mapped to the negative strand by aligning their reverse complementary sequences. We only retained the reads aligned to the concatenated *rpoB* and *rpoC* region, whose 1-based coordinate is [227354, 235682]. We then pooled the aligned reads to construct the testing data. To evaluate accuracy of iGDA, we downloaded assembly for each sample in the pooled data (Table S2) from NCBI (https://www.ncbi.nlm.nih.gov/assembly), and aligned the assembled genomes to the reference genome using MUMmer^36^. The union of the SNVs reported by MUMmer were used as benchmark to evaluate accuracy of detecting SNVs. The genome sequence of an individual sample is defined as a real contig, and was used to evaluate the accuracy of contigs reported by iGDA. We used the same method in the previous section to merge identical samples and obtain the relative abundance of each sample. The relative abundances range from 0.20% to 9.30%, and the average relative abundance is 3.20%.

Due to the unique sequencing mechanism of ONT, DNA methylation can affect the raw sequencing signal and substantially increase the base-calling error rate of methylated bases (Figure S3). The base caller used in the public ONT data in this study is Albacore (version 2.0) (https://github.com/Albacore/albacore). To avoid the impact of DNA methylation, we developed an algorithm to identify DNA methylation motifs in bacteria without using raw-signal of ONT data (details are in Methods). We masked loci within 5 bases to the DNA methylation motifs before applying iGDA to this dataset.

The result shows that the RSM algorithm substantially outperforms the single-locus methods to detect minor SNVs, and achieves a high accuracy (Figure 4C). With DNA methylation and bases with QV lower than 10 masked, iGDA detected 92.8% of the real SNVs at FDR lower than 1%. With masking no DNA methylation but masking bases with QV lower than 10, iGDA detected 41.3% of the real SNVs at FDR lower than 1%. Thus, masking DNA methylation increases the accuracy of the RSM algorithm (Figure 4D), which demonstrates the importance of removing DNA methylation or applying a methylation-aware base caller to detecting minor SNVs from ONT data. Masking bases with low QV can substantially increase the accuracy and different thresholds have similar accuracies (Figure 4C and Figure 4D). In contrast to PacBio data, correcting sequence context does not significantly increase detection accuracy of the single-locus methods. We speculate that this is because the prediction power of sequence context on the ONT data is weaker than that on the PacBio data (Figure 3).

DNA methylation has a large impact on the accuracy of phasing minor SNVs. With masking loci affected by methylation and bases with QV lower than 10, the average accuracy of assembled contigs is 90.7% (Figure 5A). However, without masking loci affected by methylation, the average accuracy of assembled contigs is only 54.5% (Figure 5A). An IGV snapshot of methylation-masked contigs is shown in Figure S4. The result shows that the iGDA-inferred contigs match the real contigs very well with DNA methylation masked. It is critical to reduce the impact of DNA methylation by whole genome amplification or by adopting a methylation-aware base caller.

### *De novo* identification of multiple *Borrelia burgdorferi* strains from long-read metagenomic data

To test whether iGDA can be applied to identify multiple strains of the same species from metagenomic data, we constructed a metagenomic dataset by mixing PacBio sequencing data of 11 *Borrelia burgdorferi* strains, the causal agent of Lyme disease^37^, and 744 other bacterial samples. The SRA IDs, species, and strains are in Table S3. We filtered the reads by requiring read quality value greater than 0.75. Read quality (rq) was extracted from the native HDF files. Bases with QV less than 8 were masked. We then aligned the reads to the reference genome of *Borrelia burgdorferi* B31 (NCBI reference sequence ID is NC_001318.1), and realigned the reverse complementary of the reads mapped to the negative strand. To evaluate accuracy of iGDA, we assembled genome of each *Borrelia burgdorferi* strain using flye^38^, and aligned the assembly to the reference genome using MUMmer^36^ to obtain benchmark SNVs.

We ran iGDA on the realigned data and constructed 1,151 contigs. The average accuracy of the contigs is 95.0% (Figure 5A) and contig length is up to 139 kb. The IGV snapshots of the contigs reported by iGDA show that multiple strains of *Borrelia burgdorferi* can be clearly identified by iGDA (Figure 6A, Figure S5, and Figure S6). The minimal divergence of a region where the *Borrelia burgdorferi* strains can be distinguished is 0.011% (details are in Methods). To further evaluate the accuracy of iGDA, we performed MLST (Multilocus Sequence Typing)^39^ on the contigs and the genome sequence of each strain using the database at https://pubmlst.org/borrelia (details are in Methods). In MLST, we aligned iGDA-inferred contigs and the genome sequence of each strain to the MLST database, consisting of known alleles of the eight house-keeping genes in *Borrelia* spp., to find the best matches. The result shows that most of the alleles that present in the genome sequence of each strain can be found in the iGDA-inferred contigs, and there is no false positive alleles (Figure 6B). The alleles of the adjacent house-keeping genes, pyrG, recG, clpX, and pepX, can be phased by the contigs reported by iGDA (Figure 6B).

**Figure 6:**
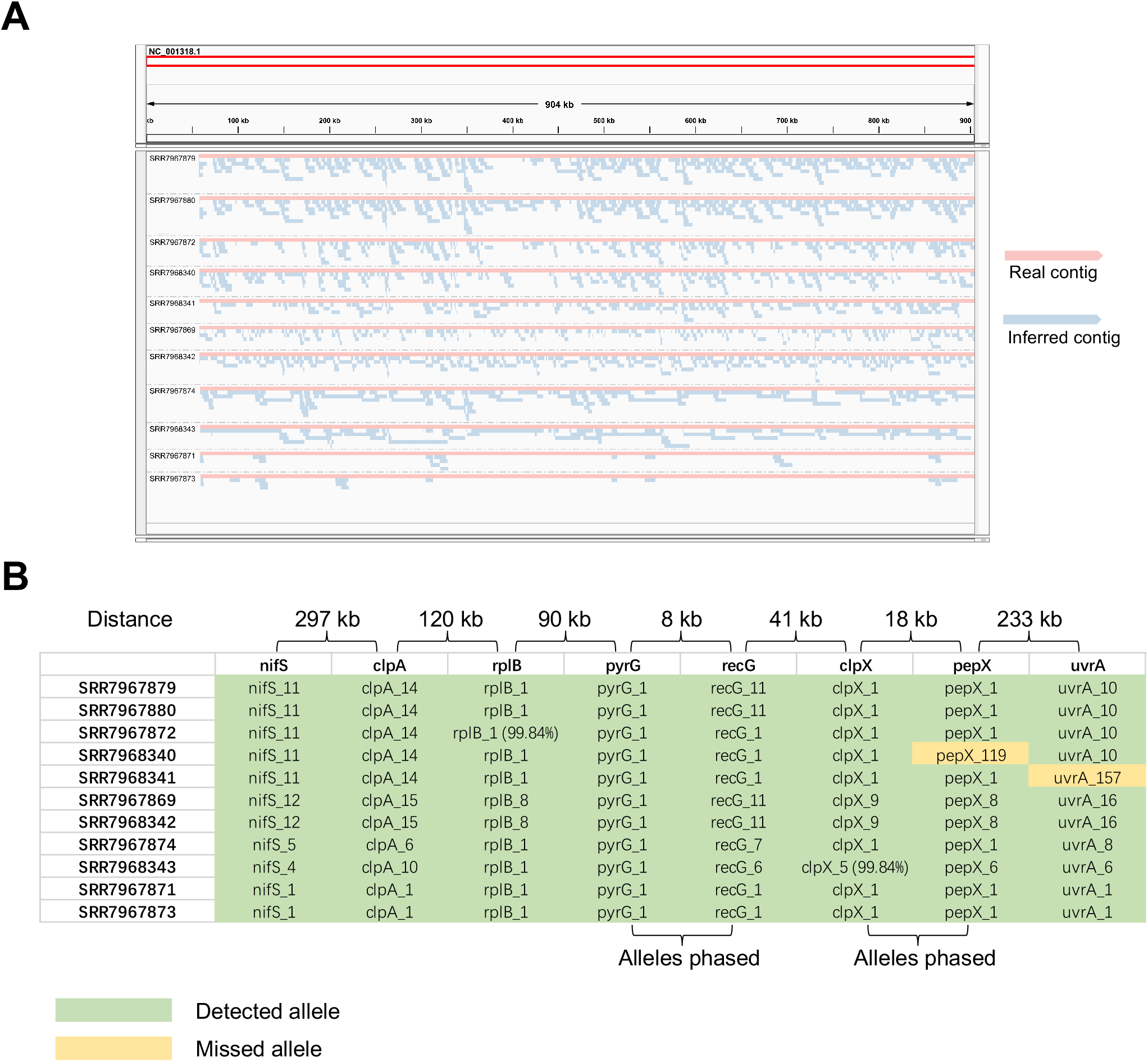
*De novo* identification of multiple *Borrelia burgdorferi* strains from PacBio metagenomic data. **A**, The IGV snapshot of the contigs inferred by iGDA from the metagenomic data. Each contig is grouped with its closest real contig (*B. burgdorferi* strain). **B**, MLST of *B. burgdorferi* in the metagenomic data. The columns are the alleles of the 8 house-keeping genes used in MLST. Each row is the alleles of the genome of each sample (strain). The row names are the SRA IDs of each sample. An allele is detected if it matches a contig inferred by iGDA. There are two alleles that have no 100% match in the MLST database, and their similarities to the closest alleles in the database are shown in the brackets. All the other alleles have a 100% match in the database.

It is worth to note that some genome regions in Figure 6A are not covered by any contig. We call these regions missed regions, and call the SNVs not covered by any contig missed SNVs. We found that there are usually multiple strains that are highly similar to each other in the missed region. In the example shown in Figure S5, at least four samples have highly similar sequences in the missed region. Some missed regions have no SNV compared to the reference genome because iGDA does not report contigs with no SNV. In the example in Figure S6, samples SRR7967871 and SRR7967873 have several large missed regions, which have no SNV compared to the reference genome. To further assess the impact of highly similar strains on the performance of iGDA, we calculated Jaccard index of SNVs for each pair of the *Borrelia burgdorferi* samples, and found that some samples are highly similar to each other. The result in Figure S7A indicates that samples SRR7967879, SRR7967880, SRR7967872, SRR7968340 and SRR7968341 are highly similar to each other, and sample SRR7967869 is highly similar to sample SRR7968342. We constructed a new dataset where only one sample is retained out of the highly similar strains. Specifically, we excluded samples SRR7967879, SRR7967880, SRR7967872, SRR7968340 and SRR7968342 from the samples listed in Table S3, and reran iGDA on the new data. The result shows that the accuracy of each contig is not significantly changed by excluding highly similar strains (Figure S7B). However, the length of contigs and proportion of SNVs covered by contigs are substantially increased (Figure S7C and Figure S7D). The species other than *Borrelia burgdorferi* have limited impact on the results, because most of the reads from these species (Table S3) cannot be aligned to the reference genome of *Borrelia burgdorferi*, and 99.93% of the aligned reads are aligned to 16S ribosomal RNA or 23S ribosomal RNA.

## Discussion

We here present iGDA, a novel open-source tool implementing several innovative algorithms that can achieve a high accuracy for detecting and phasing minor SNVs. iGDA makes it feasible to study a number of previously challenging problems, such as constructing strain-level genome sequence in microbiome samples, and identifying genome sequence of pathogens in samples with co-infection. The RSM and ANN algorithms proposed in this work are generic methods and can be extended to apply to single-cell genome sequencing data or 10X genomics linked-read^40^ data. In addition to genome sequencing, these algorithms have the potential to be applied in RNA sequencing data as well. For example, with an alternative prepossessing procedure, these algorithms can be used to decipher the heterogeneity of A-to-I RNA editing using long-read sequencing.

A major limitation of iGDA is that its high accuracy relies on the presence of multiple SNVs. Therefore, iGDA has reduced accuracy to detect orphan SNVs with very low frequency. Besides, presence of highly similar genomes will reduce accuracy of iGDA.

DNA methylation can induce correlated substitution errors on ONT data and reduce the accuracy of iGDA. Masking DNA methylation can increase the accuracy of iGDA on ONT data. Using whole genome amplification (WGA) to remove DNA methylation is a solution to this issue. Another solution is to use a base caller that can correct methylation induced error, but there is no such tool currently available according to our best knowledge.

In this work, we only detect minor SNVs because they are less affected by alignment ambiguity compared to insertions and deletions (Indel). Alignment ambiguity means an Indel might be located to multiple loci in the genome but the corresponding alignment scores are equal. To extend our RSM and ANN algorithms to detect minor Indels or other more complicated variants, an alternative way to represent variants and alignments is needed.

## Methods

### Leveraging multiple loci to detect SNVs

For the *i*th aligned read, we encode its substitution at locus *k* of the reference genome by the following formula:

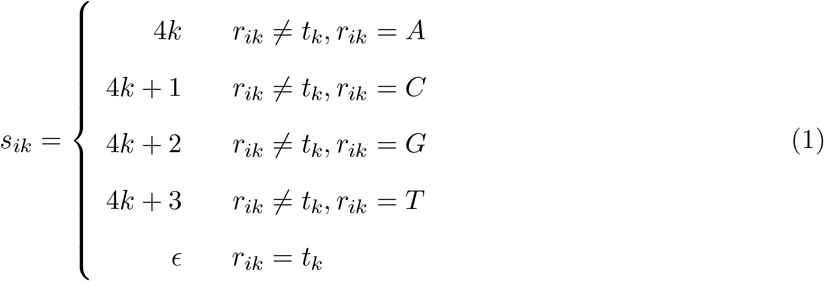

where *r*_*ik*_ is the base (short for nitrogenous base) of the *i*th aligned read at locus *k, t*_*k*_ is the base at locus *k* of the reference genome and *ϵ* is an empty element, which is formally defined by *{ϵ}* = ∅. The first locus of the reference genome is 0 throughout this paper unless otherwise stated. The *i*th read is represented as a set of substitutions and its covering range (Figure 2A), and is denoted by

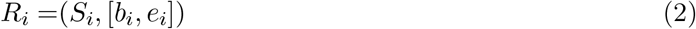

*b*_*i*_ and *e*_*i*_ are the start and end loci of the region covered by the read respectively, and *S*_*i*_ is

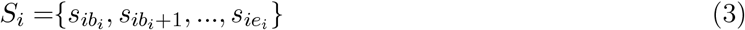

The most intuitive way to detect SNVs is to use the substitution rate of each locus. Formally, we denote the encoded substitution at locus *k* as a random variable *X*_*k*_, and denote probability of the event {*X*_*k*_ = *x*_*k*_} as *Pr*(*X*_*k*_ = *x*_*k*_), where *x*_*k*_ *∈ {4*k*, 4*k* + 1, 4*k* + 2, 4*k* + 3*}. Substitution rate is defined as the estimated *Pr*(*X*_*k*_ = *x*_*k*_), which is

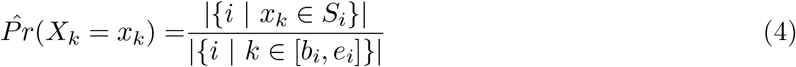

where {*·}* is a set and |*·*| is the number of elements in a set. Intuitively, in equation (4), the numerator is the number of reads with substitution *x*_*k*_ at locus *k*, and the denominator is the number of reads covering locus *k*. Due to the high error rate of long-read sequencing data, it is inaccurate to detect minor variants using substitution rate alone (Figure 1B). Herein, we leverage the information of multiple loci to increase the detection accuracy. Assuming sequencing errors are independent with each other, real SNVs are likely to be present if there are multiple reads containing the same set of substitutions (Figure 1A). The conditional probability of {*X*_*k*_ = *x*_*k*_} given other real SNVs of the same genome is therefore much larger than the marginal probability of {*X*_*k*_ = *x*_*k*_} if *x*_*k*_ is a real SNV, because these real SNVs are positively dependent (Figure 1A and Figure 1C). Formally, the conditional probability of event {*X*_*k*_ = *x*_*k*_} given *p* other substitutions is defined as 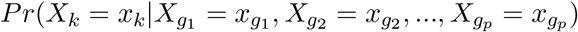, which is estimated by

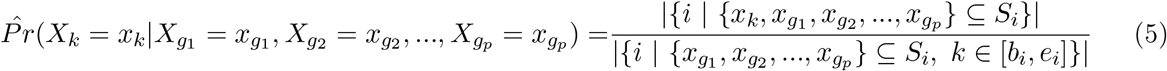

Intuitively, in equation (5), the numerator is the number of reads containing substitution *x*_*k*_ and the *p* other substitutions, and the denominator is the number of reads that contain the *p* other substitutions and cover locus k. The *p* loci, *g*_1_, *g*_2_, …, and *g*_*p*_ are called dependent loci. As 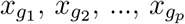, and *p* in equation (5) are unknown, the estimated maximal conditional probability of event {*X*_*k*_ = *x*_*k*_} given *p* other substitutions is used to detect SNVs, and is formally defined by

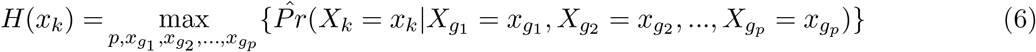

The substitution *x*_*k*_ is detected as a real SNV if *H*(*x*_*k*_) is larger than a threshold (0.65 in this study). *H*(*x*_*k*_) is also called maximal conditional substitution rate. To avoid high variance of the estimated 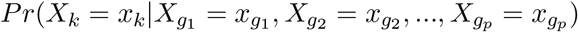 (equation (5)), we require that 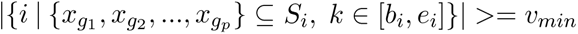, and *v*_*min*_ = 25 in this study. Sequencing errors at multiple loci that are very close to each other might induce slightly dependent substitutions. To avoid the impact of dependent substitutions induced by sequencing errors, we require that locus *k* and loci *g*_1_, *g*_2_, …,*g*_*p*_ are not too close. Specifically, we require HD(*k, g*_*s*_) *≥* 15 for any *g*_*s*_ *∈ {g*_1_, *g*_2_, *…, g*_*p*_}. HD(*k, g*_*s*_) is the homopolymer distance between locus *k* and locus *g*_*s*_, and is defined as the number of homopolymers between the two loci. A homopolymer is a set of consecutive identical bases, and a base with no identical adjacent bases is also defined as a special homopolymer with size equal to 1.

It is computationally infeasible to enumerate all combinations of *p* loci to estimate *H*(*x*_*k*_) in equation (6). It is important to note that it is insufficient to detect SNVs accurately by restricting the number of dependent loci *p* to a certain number. In the example shown in Figure S1, *H*(*x*_*k*_) fails to detect the real SNVs if *p* is restricted to 1. Likewise, we can also have similar examples if *p* is restricted to another number greater than 1. In this work, we developed a novel algorithm called Random Subspace Maximization (RSM) that can estimate *H*(*x*_*k*_) efficiently without restricting *p*.

### Detecting SNVs by RSM algorithm

#### The greedy algorithm and its theoretical accuracy

We introduce a fast but inaccurate greedy algorithm to estimate *H*(*x*_*k*_) (equation (6)), and then improve its accuracy by Random Subspace Maximization (RSM) in the next section. To estimate *H*(*x*_*k*_) for substitution *x*_*k*_ at locus *k*, we only need to consider dependent loci in range [*k* − *t*_*l*_, *k* + *t*_*r*_], where

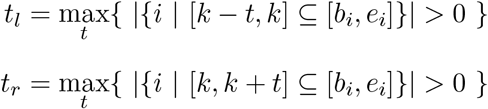

[*b*_*i*_, *e*_*i*_] is the covering range of read *R*_*i*_ (equation (2)). We estimate *Pr*(*X*_*k*_ = *x*_*k*_|*X*_*g*_ = *x*_*g*_) by equation (5) for each locus *g ∈* [*k* − *t*_*l*_, *k* + *t*_*r*_] *∩ {k}*^*c*^ ({*·}*^*c*^ is complement of a set), and sort the loci according to *Pr*(*X*_*k*_ = *x*_*k*_|*X*_*g*_ = *x*_*g*_) in descending order. The sorted loci are denoted as 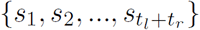, and 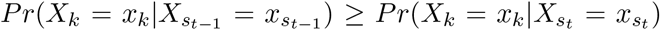. We keep adding locus *s*_*t*_ to {*s*_1_, *s*_2_, *…, s*_*t*−1_} if 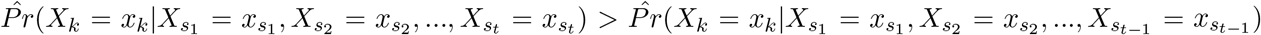 and stop if otherwise. 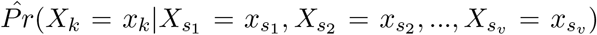 based on the final *v* selected loci {*s*_1_, *s*_2_, *…, s*_*v*_} is used to estimate *H*(*x*_*k*_).

The naive greedy algorithm described above avoids combinatorial explosion but might have low accuracy. We assume 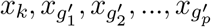 are *p* + 1 real SNVs of the same genome, and 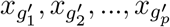 are the only *p* substitutions that can maximize 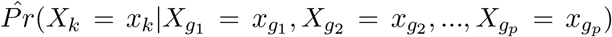. Formally, 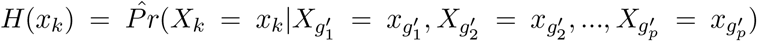, and 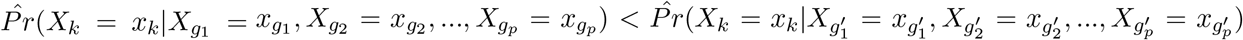 if 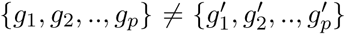. Assuming *k*, 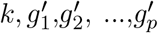 are the only loci with real SNVs in [*k* − *t*_*l*_, *k* + *t*_*r*_], we define signal-to-noise ratio by

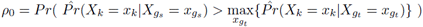

where 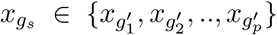 and 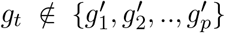. 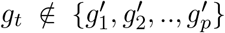 is equivalent to 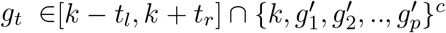. For any locus 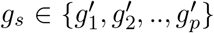, the probability that it is selected by the greedy algorithm is denoted as *Pr*(*g*_*s*_ *∈ {s*_1_, *s*_2_, *…, s*_*v*_}), where {*s*_1_, *s*_2_, *…, s*_*v*_}) is the *v* loci selected by the greedy algorithm. Without loss of generality, assuming *v* ≤ *p* and sequencing errors are independent,

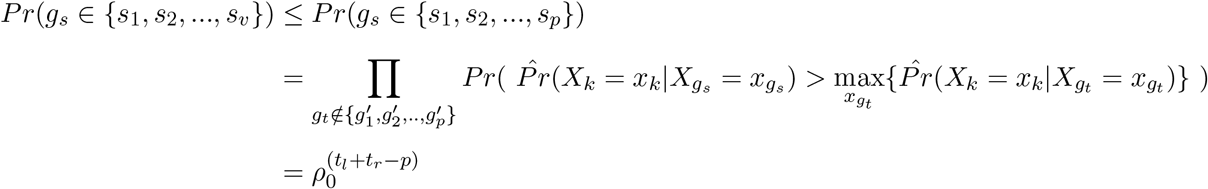

The probability that the greedy algorithm correctly estimates *H*(*x*_*k*_) is

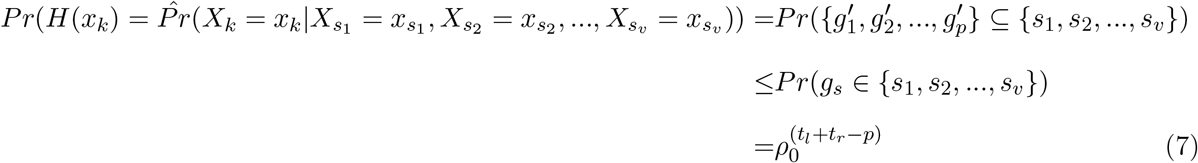

According to inequation (7), assuming *t*_*l*_ *≥* 2000, *t*_*r*_ *≥* 2000, and *p* = 1, which is a typical setting for long-read sequencing data, the probability that the greedy algorithm correctly estimates *H*(*x*_*k*_) is less than 3.5 *×* 10^−18^ even if *ρ*_0_ = 0.99. The key factor leading to the failure of the greedy algorithm is selecting from too many loci (*t*_*l*_ + *t*_*r*_ loci). We propose a novel algorithm called Random Subspace Maximization (RSM) to reduce the number of loci to be considered in the next section.

#### Improving accuracy of the greedy algorithm by Random Subspace Maximization

Firstly, we measure the similarity between two reads, *R*_*i*_ and *R*_*j*_, by a modified Jaccard index^34^, which is defined by

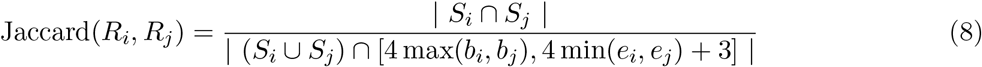

where Jaccard(*R*_*i*_, *R*_*j*_) = 0 if the denominator is 0. We require

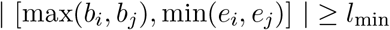

where *l*_min_ is the minimal length of the overlap region between the two compared reads. We used *l*_*min*_ = 0.5(*e*_*i*_ − *b*_*i*_) in this work. Intuitively, the Jaccard index between two reads is the ratio between number of common substitutions shared by the two reads and the total number of substitutions of the two reads in their overlapped region. Then, for a read *R*_*i*_, we select *w* most similar reads according to Jaccard index. For each read *R*_*j*_ in these *w* selected reads, we generate a set of substitutions shared by *R*_*i*_ and *R*_*j*_. Formally,

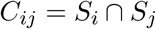

*C*_*ij*_ is called a subspace (Figure 2B), and we can generate *w × m* subspaces if there are *m* reads. We used *w* = 100 in this work. For a substitution *X*_*k*_ *∈ C*_*ij*_, we estimate its maximal conditional probability of {*X*_*k*_ = *x*_*k*_} in subspace *C*_*ij*_, which is defined by

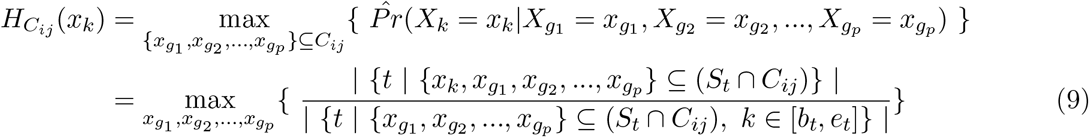

using the greedy algorithm described in the previous section by only considering the substitutions in *C*_*ij*_. Thus, compared to the original greedy algorithm, the number of loci to be considered is substantially reduced. We then use

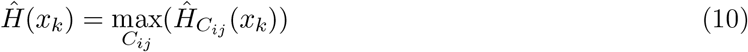

to estimate the maximal conditional probability of {*X*_*k*_ = *x*_*k*_} defined by equation (6). 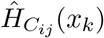 is the maximal conditional probability of {*X*_*k*_ = *x*_*k*_} in subspace *C*_*ij*_ estimated by the greedy algorithm. The whole procedure of estimating *H*(*x*_*k*_) in the *w × m* subspaces is called Random Subspace Maximization (RSM) (Figure 2C).

#### Theoretical accuracy of RSM algorithm

Without loss of generality, we denote 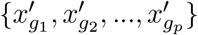 as the only set of substitutions that maximizes 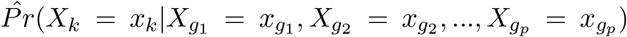, and Ω as the set of subspaces containing 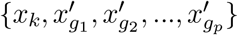. For a subspace *C*_*t*_ *∈* Ω, the probability that the greedy algorithm finds 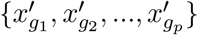 is denoted as 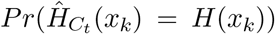, where *H*_*k*_ is defined by equation (6). The probability that RSM algorithm finds 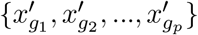 is

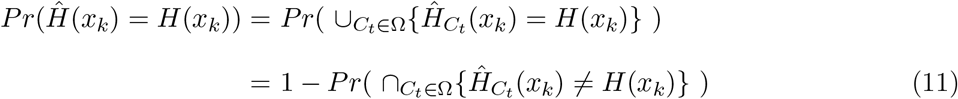

Assuming 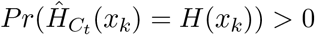, and according to chain rule of joint probability,

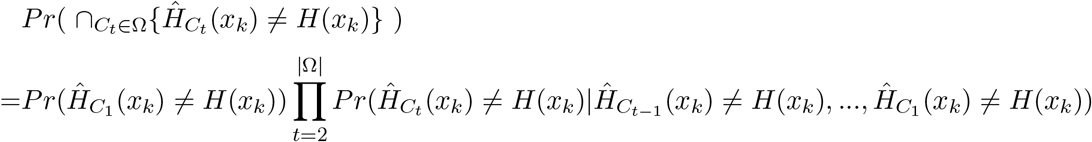

where 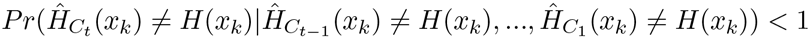 if 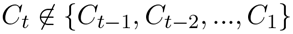. As sequencing depth increases, |Ω| increases, and 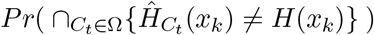 converges to 0. Thus, *Pr*(*Ĥ*(*x*_*k*_) = *H*(*x*_*k*_)) (equation (11)) converges to 1 as sequencing depth increases. Intuitively, with infinite sequencing depth, RSM algorithm is guaranteed to detect real SNVs correctly if these SNVs have larger maximal conditional probabilities than sequencing errors.

#### Detecting orphan SNVs by correcting sequence context effect

As RSM algorithm requires multiple real SNVs, it can not detect orphan SNVs. An orphan SNV is the only SNV of the genome. We have to rely on the single-locus algorithm described in equation (4) to detect orphan SNVs. However, the substitution rate of a locus is not only affected by real SNVs but also affected by the sequence context of the locus. We built a gradient boosting^31^ model to learn the sequence context effect and corrected it by the following likelihood ratio method (Figure 2D). For a substitution *x*_*k*_ at locus *k*, its likelihood ratio is

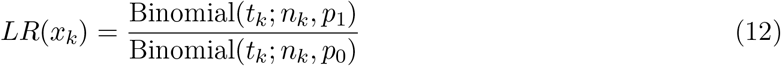

where Binomial(*x*; *n, p*) is the probability mass function of binomial distribution with parameters *n* and *p*, and

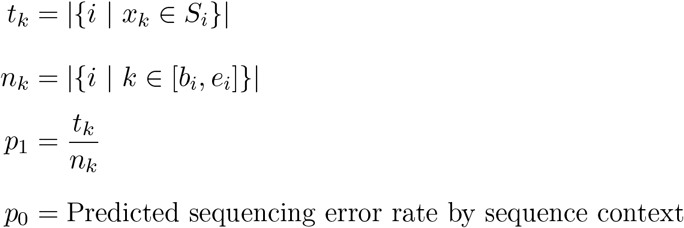

The substitution *x*_*k*_ is detected as a SNV if *LR*(*x*_*k*_) is larger than a threshold. We used a threshold of 50 in this work. Calculation of *p*_0_ is introduced in the next section. To reduce false discovery rate, we also required a detected SNV has a substitution rate higher than 0.1 for PacBio data and 0.2 for ONT data respectively.

#### Modeling sequence context effect on sequencing error rate

Error rate of long-read sequencing is strongly affected by sequence context (Figure 3). For locus *i*, we define its one upstream homopolymer and one downstream homopolymer as its sequence context (Figure S8). We adopted the gradient boosting model implemented by xgboost (version 0.90)^31^ to predict substitution rate of each locus by its sequence context. For PacBio, we trained the model on a dataset consisting of 79 PacBio RS II runs with P6-C4 chemistry and a dataset consisting of 24 PacBio RS II runs with P4-C2 chemistry respectively (SRA IDs of the data are listed in Table S4). As the sequence context effects on these two datasets are highly similar, we only used the model trained on the P6-C4 data for the analysis. For ONT, we trained the model on a dataset consisting of 8 MinION runs with R9.4 chemistry (SRA IDs of the data are listed in Table S4). We tuned three parameters in gradient boosting, step size (eta in xgboost), number of trees (num_round in xgboost) and maximal depth of trees (max_depth in xgboost) and used the parameters with the highest five-fold cross-validation accuracy (Table S5). We used *R*^2^ as the measurement of accuracy, which is defined by

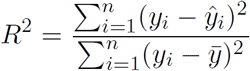

where *y*_*i*_ is the substitution rate of a sequence context, ŷ_*i*_ is the predicted substitution rate, 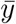 is the average substitution rate, and *n* is the number of unique sequence contexts. For PacBio, step size, number of trees and maximal depth of trees with the highest accuracy are 0.01, 2000 and 10 respectively. For ONT, step size, number of trees and maximal depth of trees with the highest accuracy are 0.1, 2000 and 10 respectively.

We also masked bases with QV thresholds 8, 10 and 12, and trained three different models on the masked data. Each model is used in the detection algorithm which masks bases with the same QV threshold. In the case of not masking any base, we predicted substitution rate using the trained model on the three pooled sequencing datasets (Figure 3). The results show that substitution-error rate is strongly affected by sequence context and can be well predicted by our model.

### Phasing minor SNVs

To detect whether multiple minor SNVs are from the same DNA molecule, we proposed a novel algorithm called Adaptive Nearest-Neighbors clustering (ANN). As the reads inevitably have errors, an intuitive way to phase minor SNVs is to cluster the reads and use the consensus sequences of each cluster to phase the minor SNVs. However, an intrinsic difficulty of clustering algorithms is to determine the number of clusters, which is unknown. The ANN algorithm can directly estimate the number of clusters from data.

#### Adaptive-Nearest-Neighbors clustering

Firstly, to reduce noise level, we only retain detected SNVs for each read. Formally, for read *R*_*i*_ (equation (2)), we use

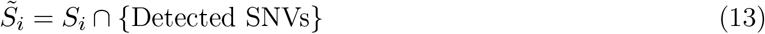

where *S*_*i*_ is defined in equation (3).

The intuitive idea of ANN algorithm is that all loci should be homogeneous by piling up the reads in each cluster (Figure S9). A locus is homogeneous if it satisfies the following condition. For locus *k*, its substitution rate satisfies

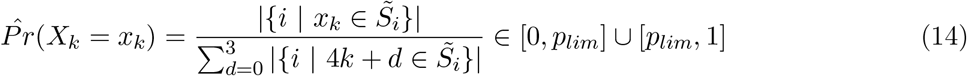

where *x*_*k*_ *∈ {*4*k*, 4*k* + 1, 4*k* + 2, 4*k* + 3}. In this work, We set *p*_*lim*_ = 0.2 for the PacBio data and *p*_*lim*_ = 0.3 for the ONT data. For a read *i* (called seed read), we sorted its *q* most similar reads according to Jaccard index (equation (8)), and kept discarding the most dissimilar one until all loci covered by the seed read are homogeneous or maximal coverage of the loci is smaller than a threshold (10 in this work) (Figure 2F). We recorded the consensus sequence as a draft contig if all the loci are homogeneous (Figure S9). We calculated the Jaccard index of each read with all the draft contigs, and assigned the read to the contig with the largest Jaccard index. A read is assigned to the reference genome if its largest Jaccard index is smaller than 0.5. The abundance of a contig is defined as the number of reads assigned to it.

A problem of the algorithm described above is that the alignment is affected by reference bias and homogeneous loci could be mistaken for heterogeneous loci. Reference bias is the phenomenon that the substitution rate of a real SNV at a homogeneous locus is significantly lower than 1 − substitution error rate (Figure S10A).

#### Reference bias and local realignment

For each detected SNV, we adopted standard Smith-Waterman algorithm implemented by SeqAn (version 2.4) (https://www.seqan.de) to realign reads to four modified reference sequences with A,C,G, or T at each locus with a detected SNV. The scores of match, mismatch, gap open, and gap extension are 2, −4, −4, and −2 respectively, and the score of a base aligned to base N or a masked low-QV base is 0. To avoid high computational cost, we only realigned 21 homopolymers whose center is the locus with detected SNV. For each read, the modified base in the reference sequence with the highest alignment score is recorded as a substitution of the read (Figure 2E and Figure S11). We tested the realignment method on a single *Escherichia coli* dataset (SRA ID is ERS718594), which is presumably homogeneous. The result shows that local realignment can substantially reduce reference bias (Figure S10B). The average substitution rate of loci with real SNVs is 84.8% before realignment, and the average substitution rate of loci with real SNV is 95.9% after realignment. We performed local realignment before ANN algorithm in our analysis.

#### Filtering draft contigs

To reduce false positive rate of the inferred draft contigs by ANN algorithm, we adopted a two-step algorithm to filter the draft contigs (Figure 2G). In the first step, we tested whether the frequency of each individual SNV in each contig is significantly higher than the sequencing error rate and whether SNVs in each contig are independent using Bayes factor. The contig is filtered if the frequency of any of its SNVs is not significant and its SNVs are independent (Figure S12A). In the second step, we compared the contigs pairwise, and the contig with lower abundance in each pair is filtered if the contigs are not significantly different according to Bayes factor (Figure S12B).

#### Assembling draft contigs

The length of the draft contigs obtained by ANN algorithm is usually smaller than genome size, except in a few cases like a virus genome. Therefore, we have to assemble the draft contigs to obtain the whole picture of the underlined genomes in the sequenced sample. We borrowed the idea of overlap graph^32^ from *de novo* genome assembly to assemble the draft contigs. We denoted each draft contig as a vertex in a graph and compared the contigs pairwise. For a draft contig *i*, we linked it to another draft contig *j* by adding a edge from vertex *i* to vertex *j* if all the three criteria are met: 1) the two draft contigs are identical in their overlapped region; 2) the number of overlapped SNVs is more than 50% of the number of SNVs in contig *i* or that in contig *j*, or the length of overlapped region is more than 50% of the length of contig *i* or that of contig *j*; 3) the genome coordinate of the end locus of contig *i* is smaller than that of contig *j*. We then removed redundant edges by transitive reduction^41^ (Figure S13A and Figure S13B). A contig is constructed by concatenating draft contigs which are in an unambiguous path. A path is an unambiguous path if the three criteria are met: 1) in-degree of the start vertex is not 1; 2) out-degree of the end vertex is not 1 or a daughter vertex of the end vertex has more than one parental vertices; 3) in-degrees and out-degrees of the vertices other than the start vertex and the end vertex are 1 (Figure 2H and Figure S13C). We then filtered the contigs using the two-step filter introduced in the previous section. We calculated the Jaccard index of each read to all the contigs, and assigned the read to the contig with the largest Jaccard index. A read is assigned to the reference genome if its largest Jaccard index is smaller than 0.5.

### Detecting bacterial methylation motifs from ONT data without raw signal

As the raw-signal files of ONT data are usually huge and not publicly available, we developed an algorithm to detect DNA methylation motifs without raw signal. For each individual ONT data file before pooling, we extracted the flanking sequences (40 bp long) of loci whose substitution rates are greater than 0.15, and detected motifs in the flanking sequences using the motif caller developed by PacBio (https://github.com/PacificBiosciences/MotifMaker)^42^. We only retained the motifs that matches the known bacterial methylation motifs in REBASE (http://rebase.neb.com/rebase/rebase_methylase_recseqs.txt)^43^. Thus, our methylation-motif detection algorithm is conservative and only detects known motifs. We only discovered two known motifs, CCWGG and CGCATC, on the ONT data. W represents A or T.

### *Borrelia* MLST

We downloaded the allele sequences of the eight house-keeping genes from https://pubmlst.org/bigsdb?db=pubmlst_borrelia_seqdef&page=downloadAlleles, and aligned them to the iGDA-inferred contigs and the genome sequence of each *Borrelia burgdorferi* strain using MUMmer (version 3)^36^. If a contig or genome sequence has no 100% match in the allele database, we reported the allele with the highest percent identity in the MUMmer output.

### Evaluating the minimal divergence that two conspecific strains can be ditinguished

We only retained the iGDA-reported contigs that is 100% identical to a true genome sequence and only has an unique closest true genome sequence. These retained contigs can be used to distinguish conspecific strains. We calculated the divergence between two contigs by

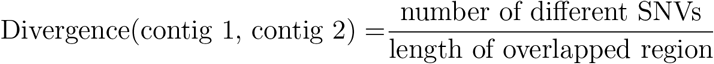

### Parameter setting in the third-party tools

#### flye

In the PacBio metagenomic data, we used “flye -t 16 –pacbio-raw -g 2m”.

#### MUMmer

We used “nucmer -c 150 -g 500 -l 12 –maxmatch” for alignment, and “show-snps -l -T -H” to obtain SNVs. To avoid the impact of repeats we used “mummerplot −−filter” before “show-snps -l -T -H” for the metagenomic data.

### Software access

iGDA is available at Anaconda Cloud https://anaconda.org/zhixingfeng/igda. Install Conda (https://docs.conda.io/projects/conda/en/latest/user-guide/install/) and type “conda in-stall -c zhixingfeng igda” to install iGDA and its dependencies. After installation, type “igda” for usage.

## Acknowledgement

The project was supported by funds from the Steven & Alexandra Cohen Foundation.

## Author contributions

Z.F. designed and implemented the computational models and algorithms of iGDA. B.W. proposed and tested the methods to improve speed of iGDA. Z.F. performed the data analysis with support from B.W.. Z.F. designed the experiments to evaluate iGDA on metagenomic data with the support from J.C. and E.E.S.. Z.F. wrote the manuscript with input from all authors.

