## Supplementary Figures for "Detecting and phasing minor single-nucleotide variants from long-read sequencing data"

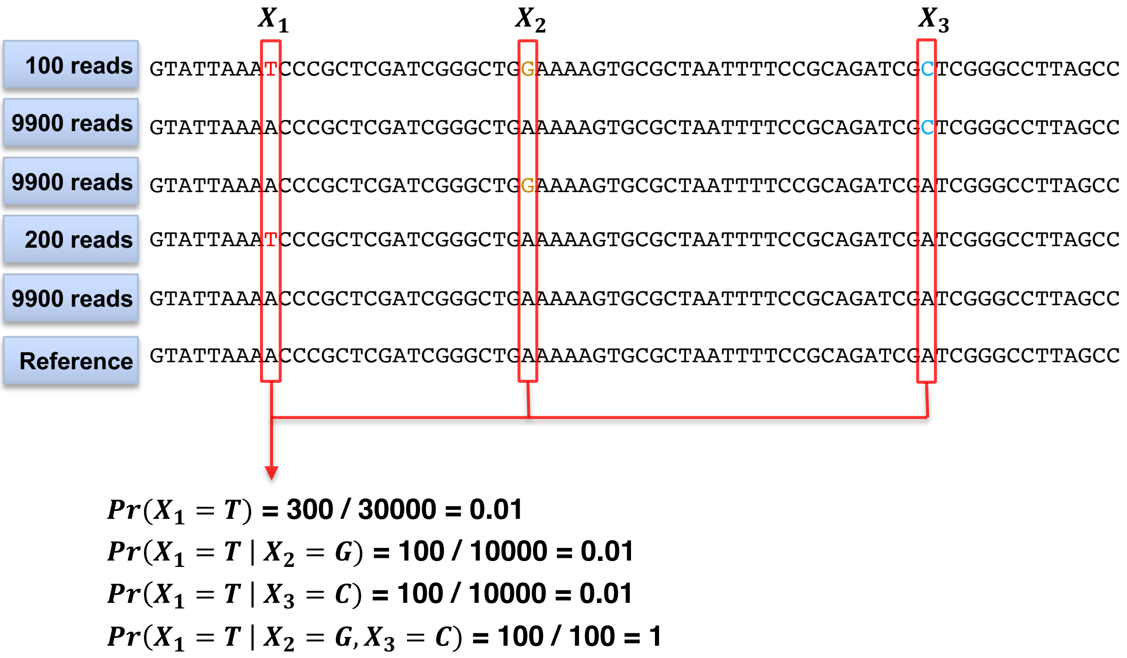


Figure S1: **Fixing the number of dependent loci might lead to false negatives.** It is difficult to detect $X_1$ by its marginal substitution rate or its conditional substitution rate given $X_2$ or $X_3$. However, $X_1$ has a large conditional substitution rate given the combination of $X_2$ and $X_3$.

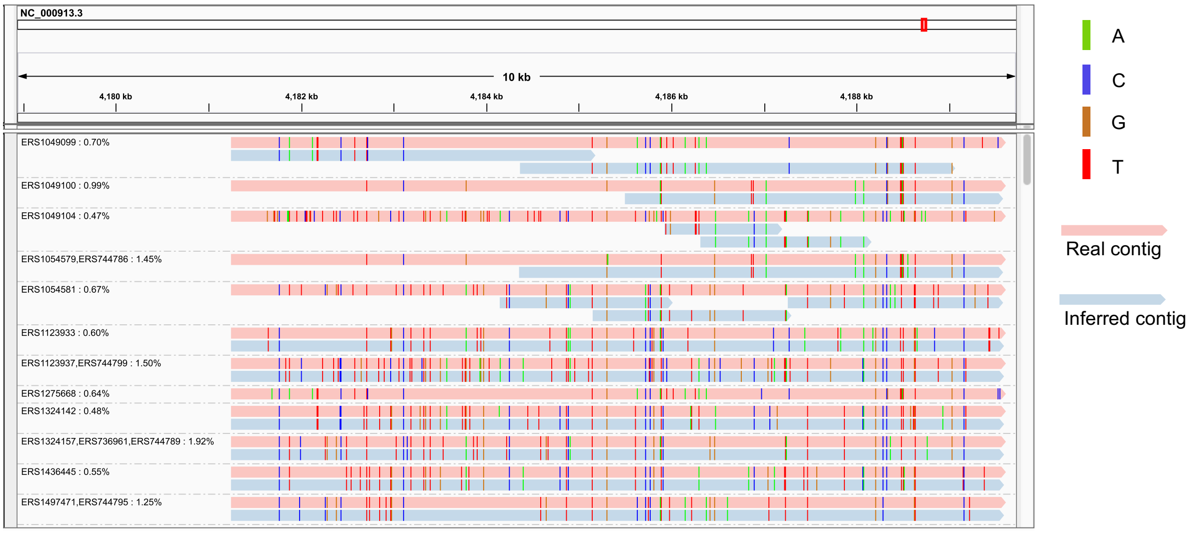


Figure S2: **The IGV snapshot of the contigs inferred by iGDA from the PacBio *E. coli* data.** Each contig is grouped with its closest real contig.

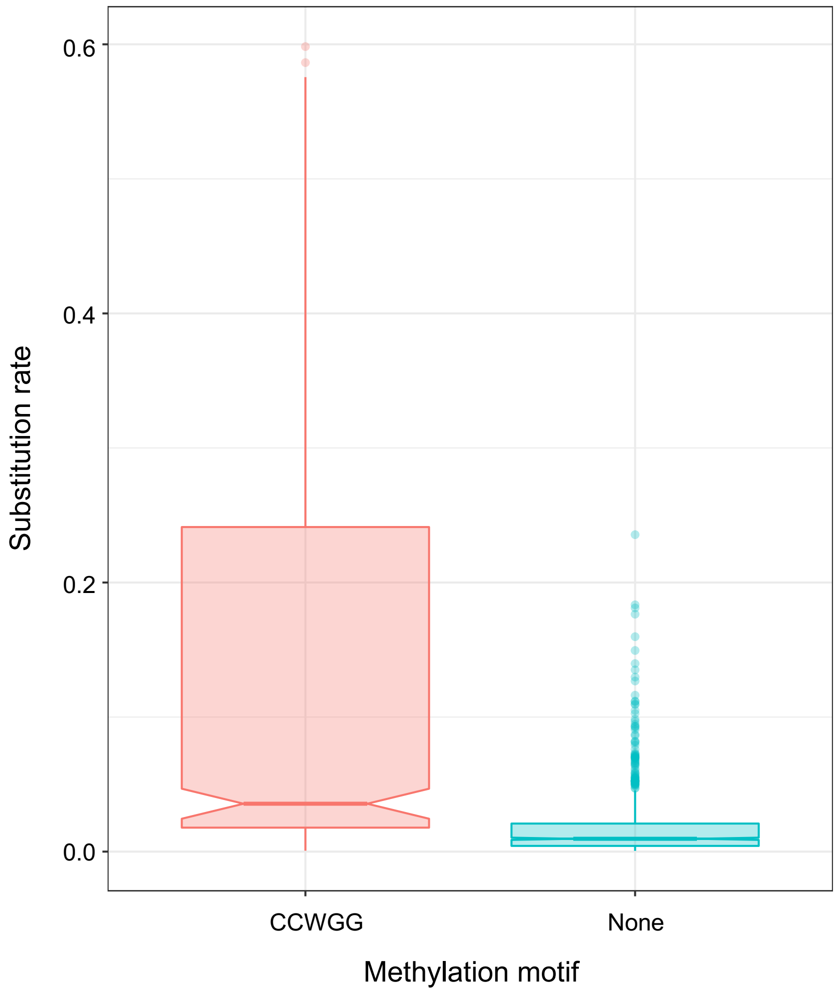


Figure S3: **Substitution rate of the loci with methylation motif.** The figure shows the substitution rate on a individual ONT dataset SRR8054586.

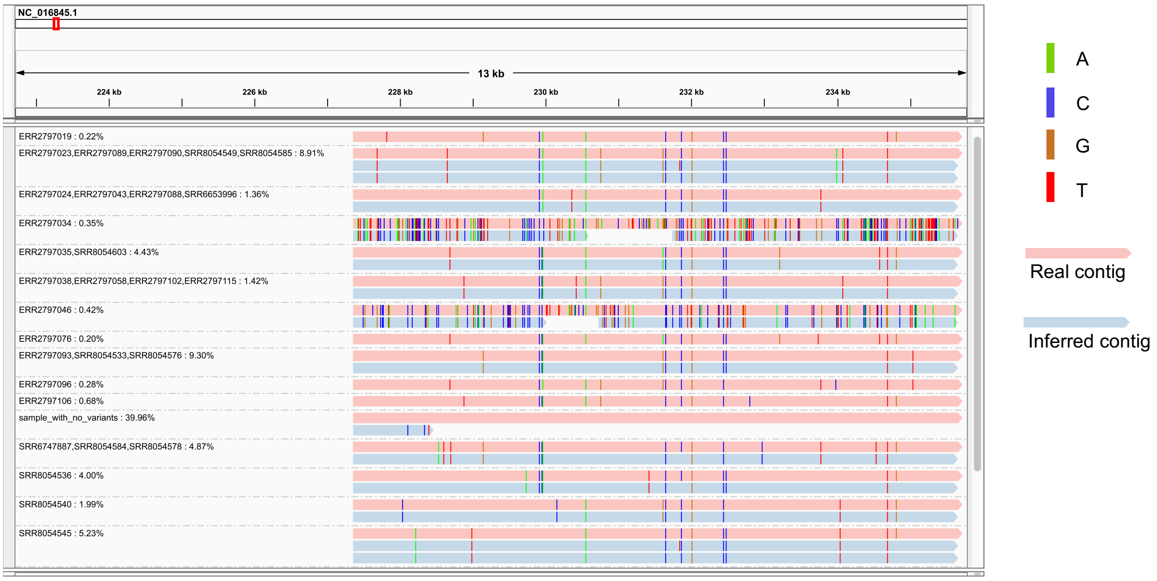


Figure S4: **The IGV snapshot of the contigs inferred by iGDA from the ONT *K. pneumoniae* data.** Each contig is grouped with its closest real contig.

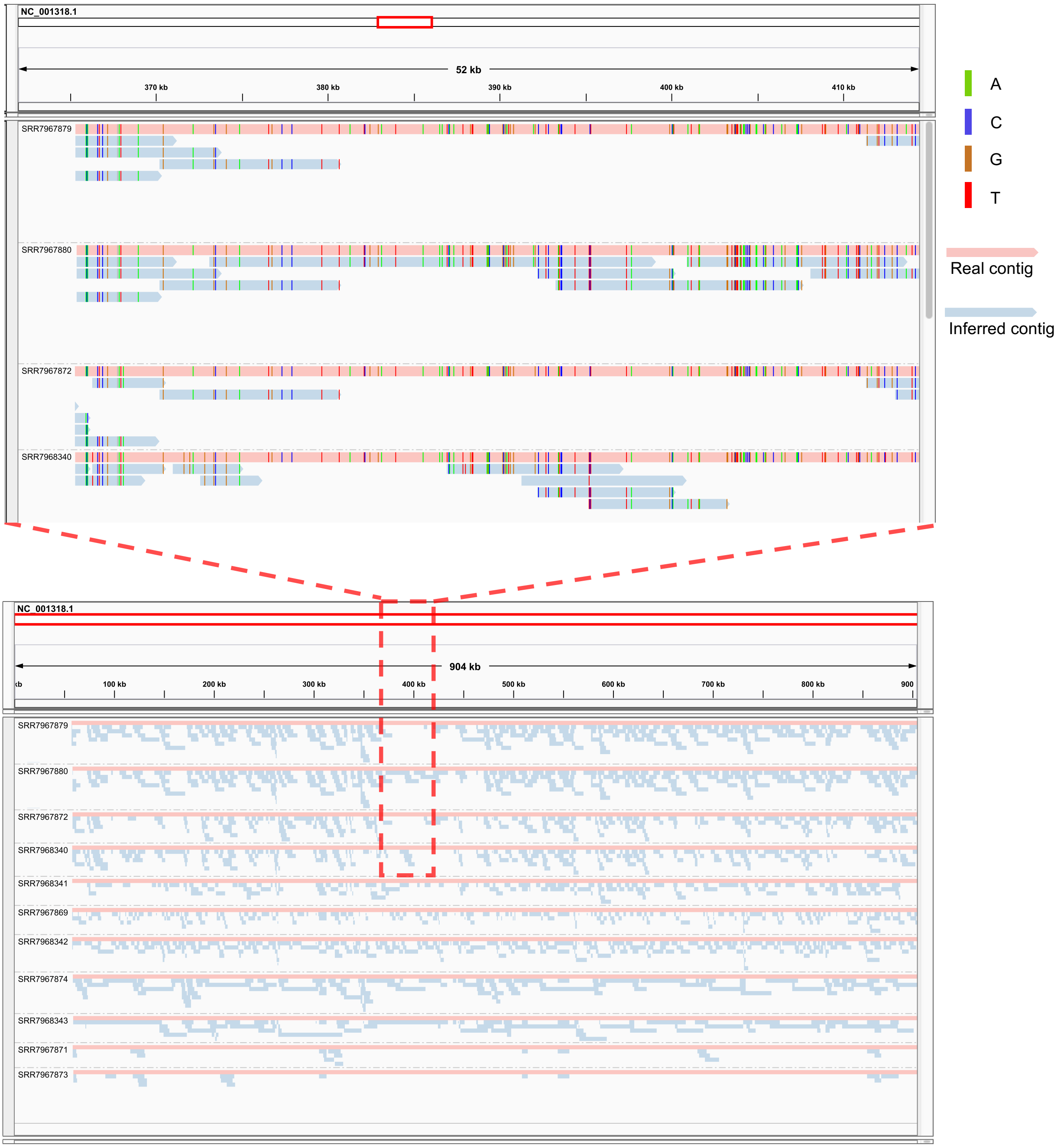


Figure S5: **Missed regions due to highly similar strains.** The lower panel shows the contigs reported by iGDA. The upper panel shows the contigs in region [363553, 449257]. Each contig is grouped with its closest real contig.

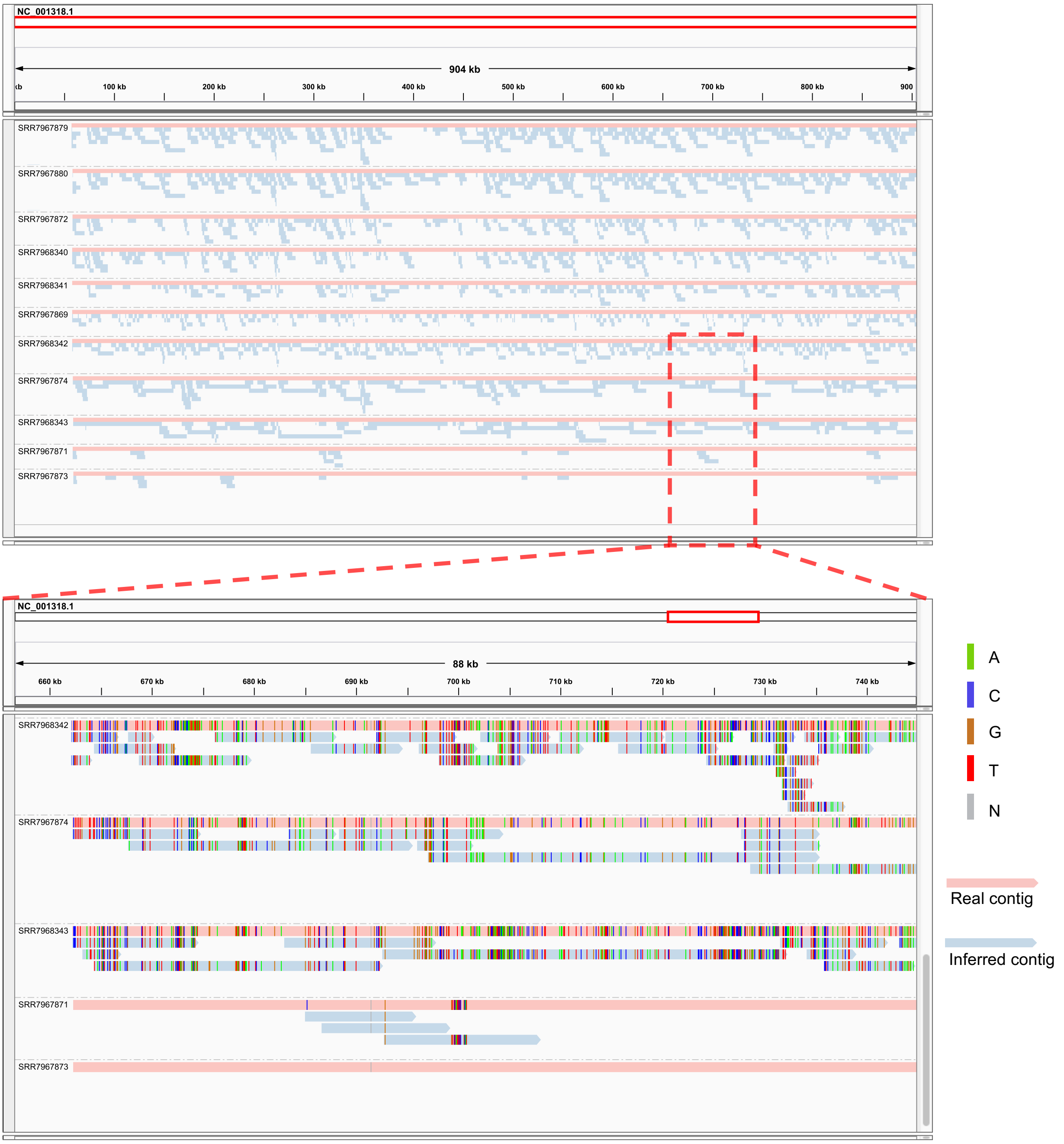


Figure S6: **Missed regions that have no SNV compared to the reference genome.** The upper panel is the contigs reported by iGDA. The lower panel is the contigs in region [656843, 745718]. Each contig is grouped with its closest real contig. Samples SRR7967871 and SRR7967873 have several large missed regions, which have no SNV compared to the reference genome.

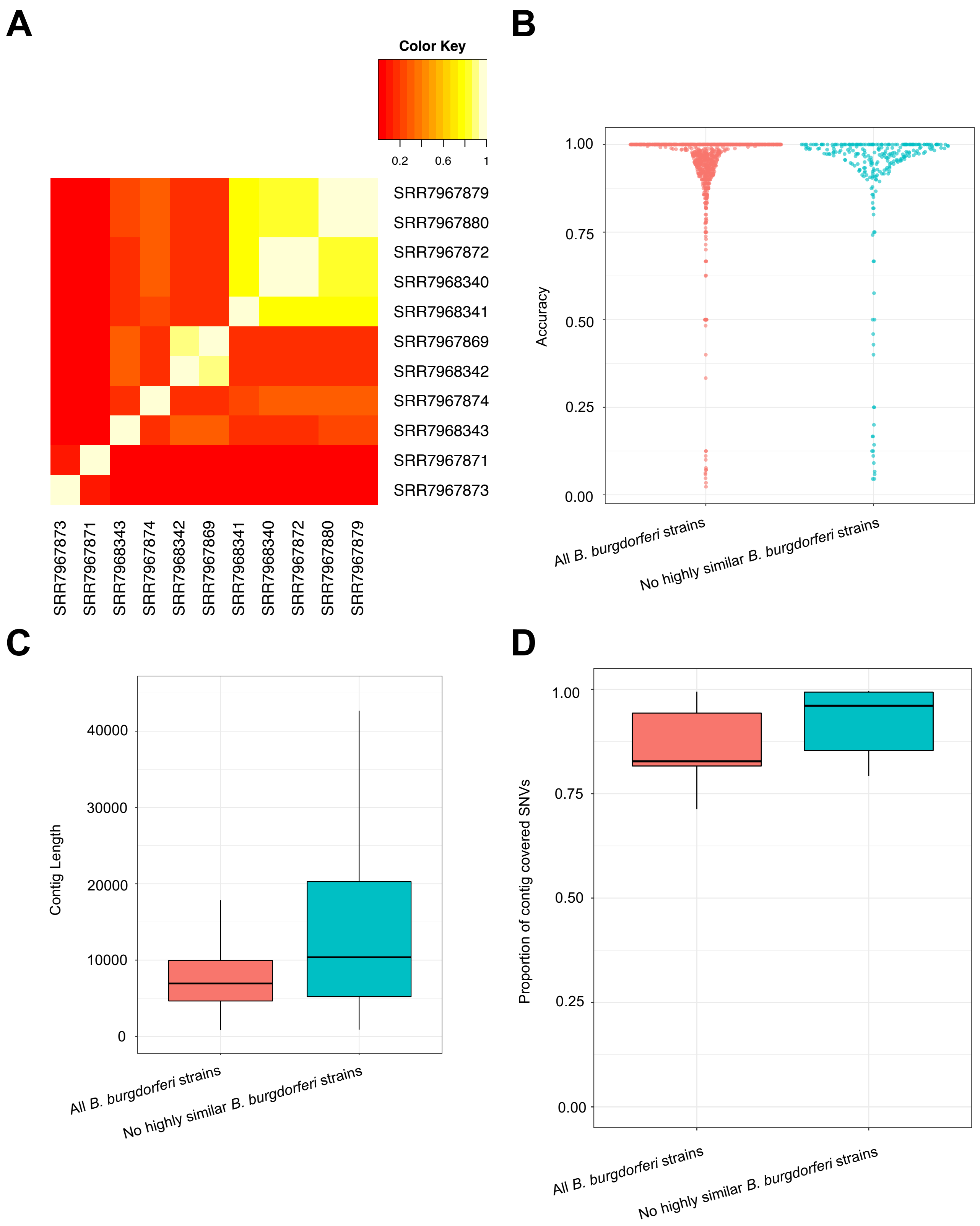


Figure S7: **Impact of highly similar strains on the performance of iGDA. A**, Heatmap of Jaccard index of the samples. **B**, Sina plot of accuracy of the iGDA inferred contigs. **C**, Box plot of contig length reported by iGDA. **D**, Box plot of proportion of SNVs covered by the contigs.

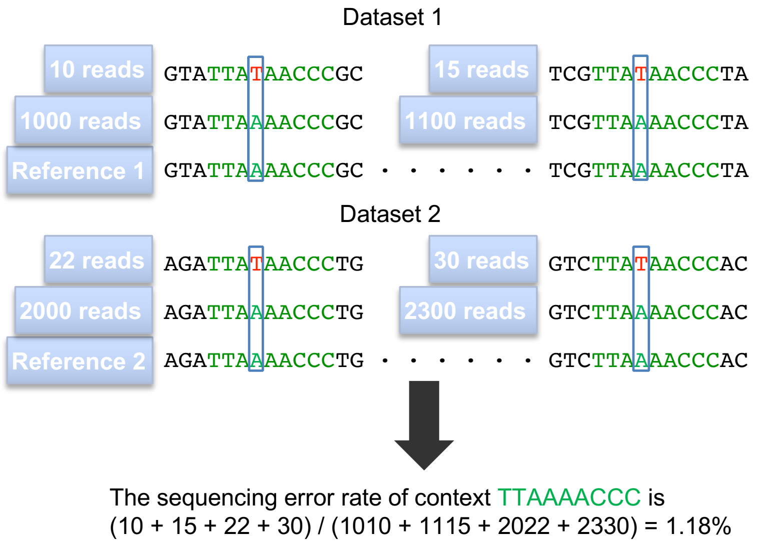


Figure S8: **Predict sequencing error rate by sequence context.** The sequence context of a locus is the concatenated sequence of one upstream homopolymer, the homopolymer containing the very locus, and one downstream homopolymer.

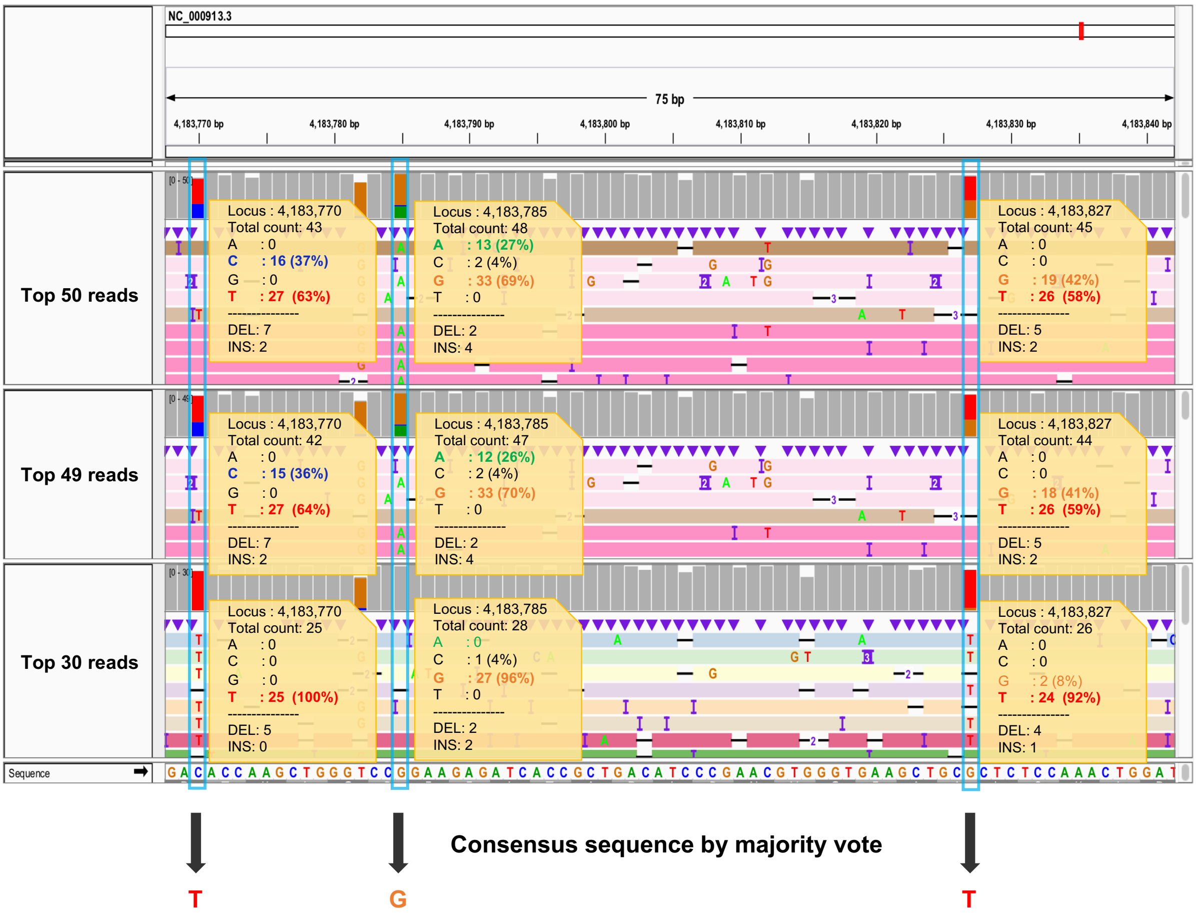


Figure S9: **An example of ANN algorithm.** Each read in the IGV snapshot is colored according to its Jaccard Index with the seed read.

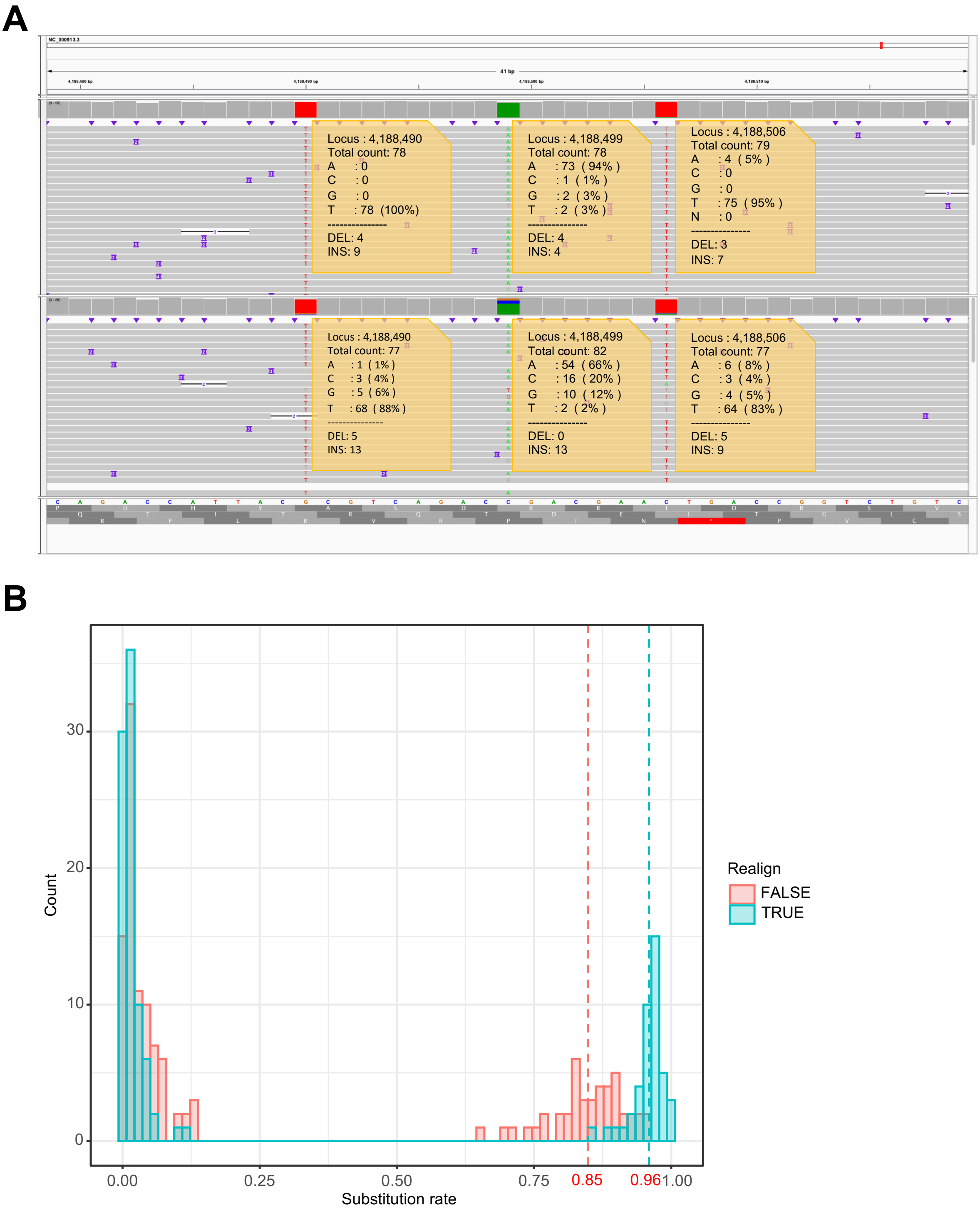


Figure S10: **Reference bias.** **A**, The IGV snapshot of PacBio sequencing data of an individual *E. coli* sample that is presumably homogeneous. The bottom track is the alignment to the original reference genome and the upper track is the alignment to the modified reference genome where the base at locus 4,188,499 is changed from C to A. **B**, The distribution of substitution rate before and after realignment.

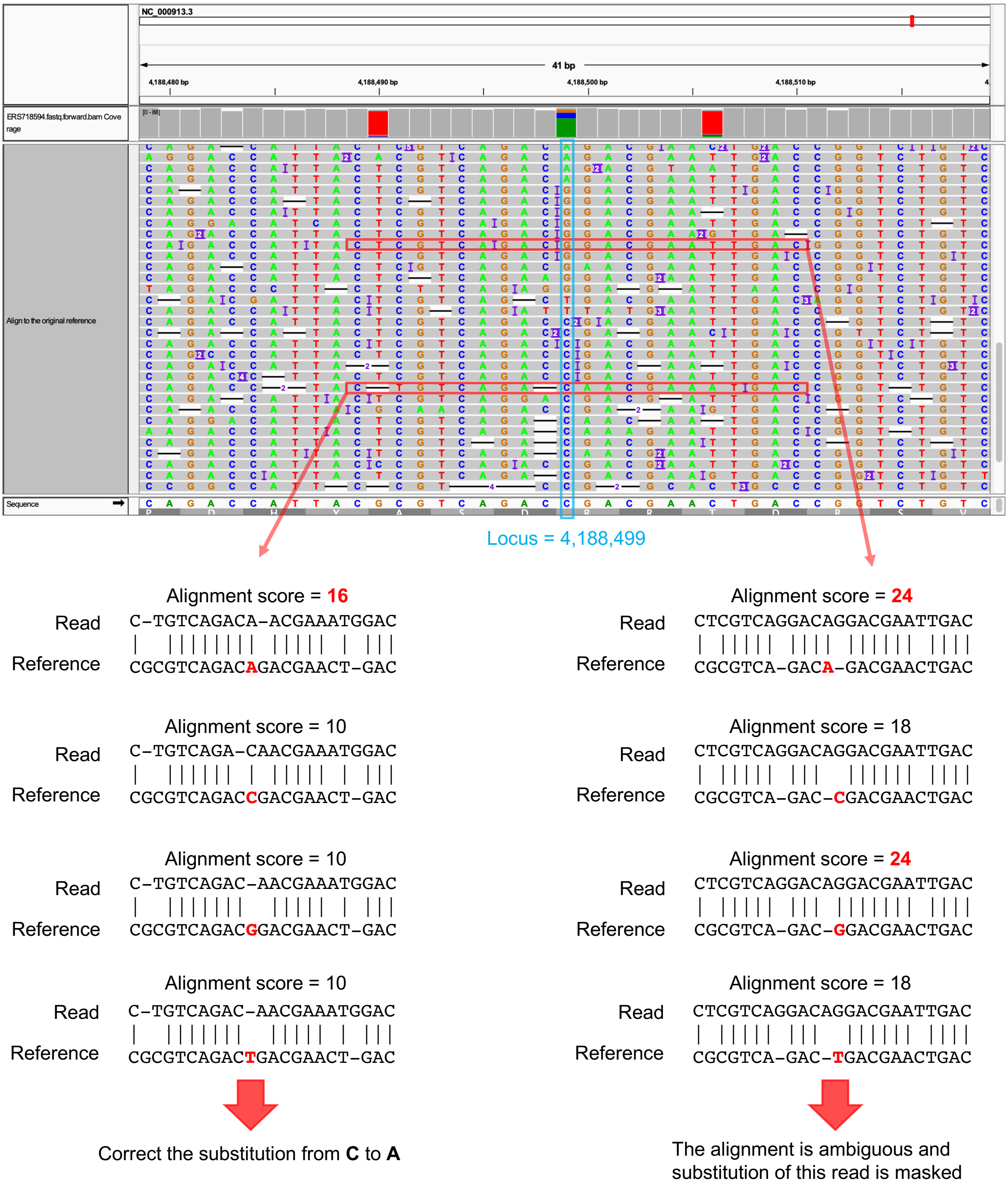


Figure S11: **Correcting reference bias by realignment.**

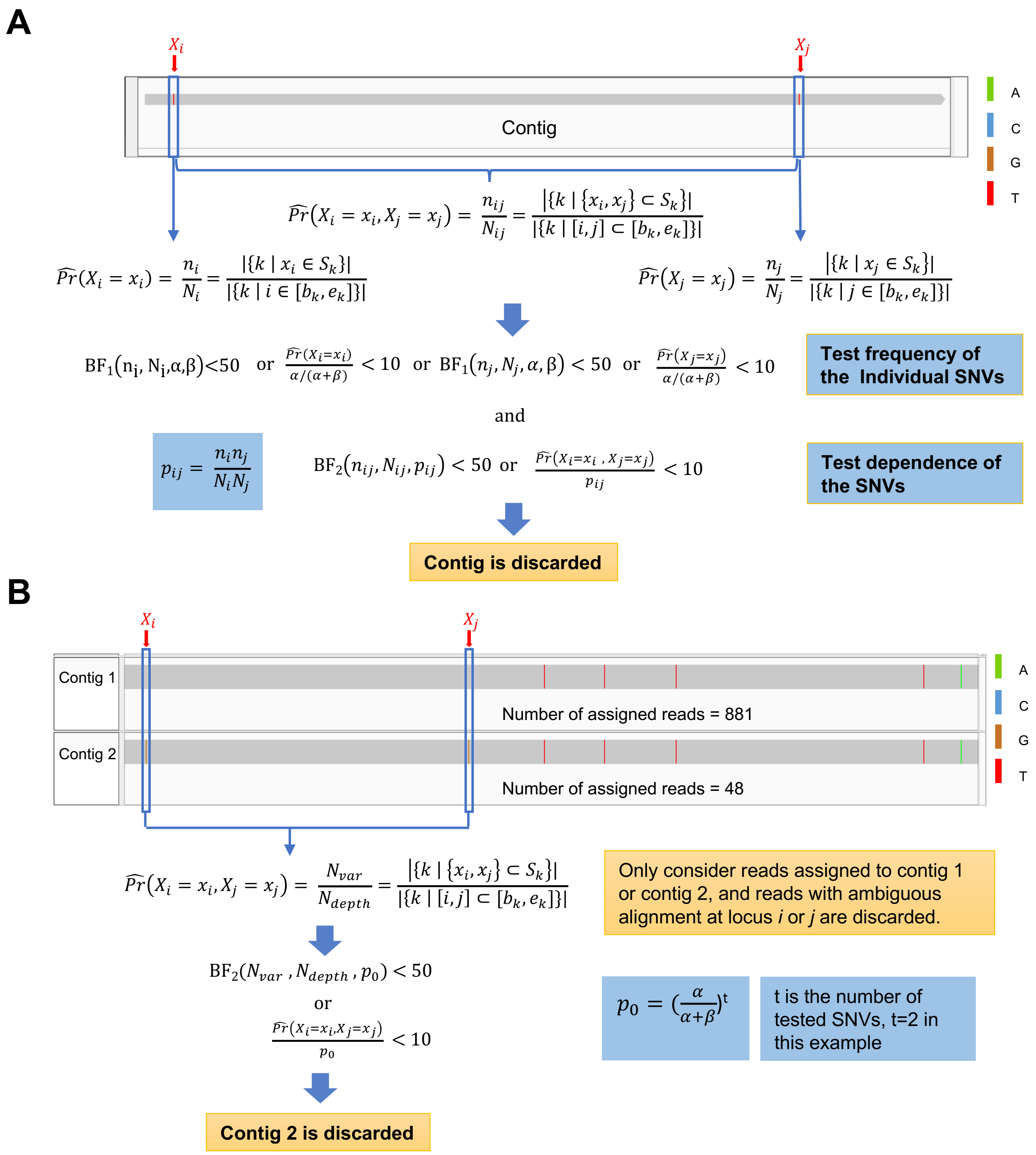


Figure S12: **Filtering contigs.** Bayes factors $\mathrm{BF}_1(n, N, \alpha, \beta) = \frac{\mathrm{Beta}(n+1, N-n+1)\mathrm{Beta}(\alpha,\beta)}{\mathrm{Beta}(n+\alpha, N-n+\beta)}$, and $\mathrm{BF}_2(n, N, p) = \frac{\mathrm{Beta}(n+1, N-n+1)}{n^p(N-n)^{(1-p)}}\left(1 - \int_0^p t^n(1-t)^{N-n}dt\right)$. $\alpha$ and $\beta$ are estimated by fitting a Beta distribution on subsitution rate obtained from the independent data in Table S4. $\alpha = 1.332824, \beta = 89.04769$ for PacBio data, and $\alpha = 0.646625, \beta = 21.90139$ for ONT data. **A**, Discarding a contig if the frequencies of all its SNVs are not significantly higher than error rate and its SNVs are independent. **B**, Discarding a contig if it is not significantly different from its similar contig.

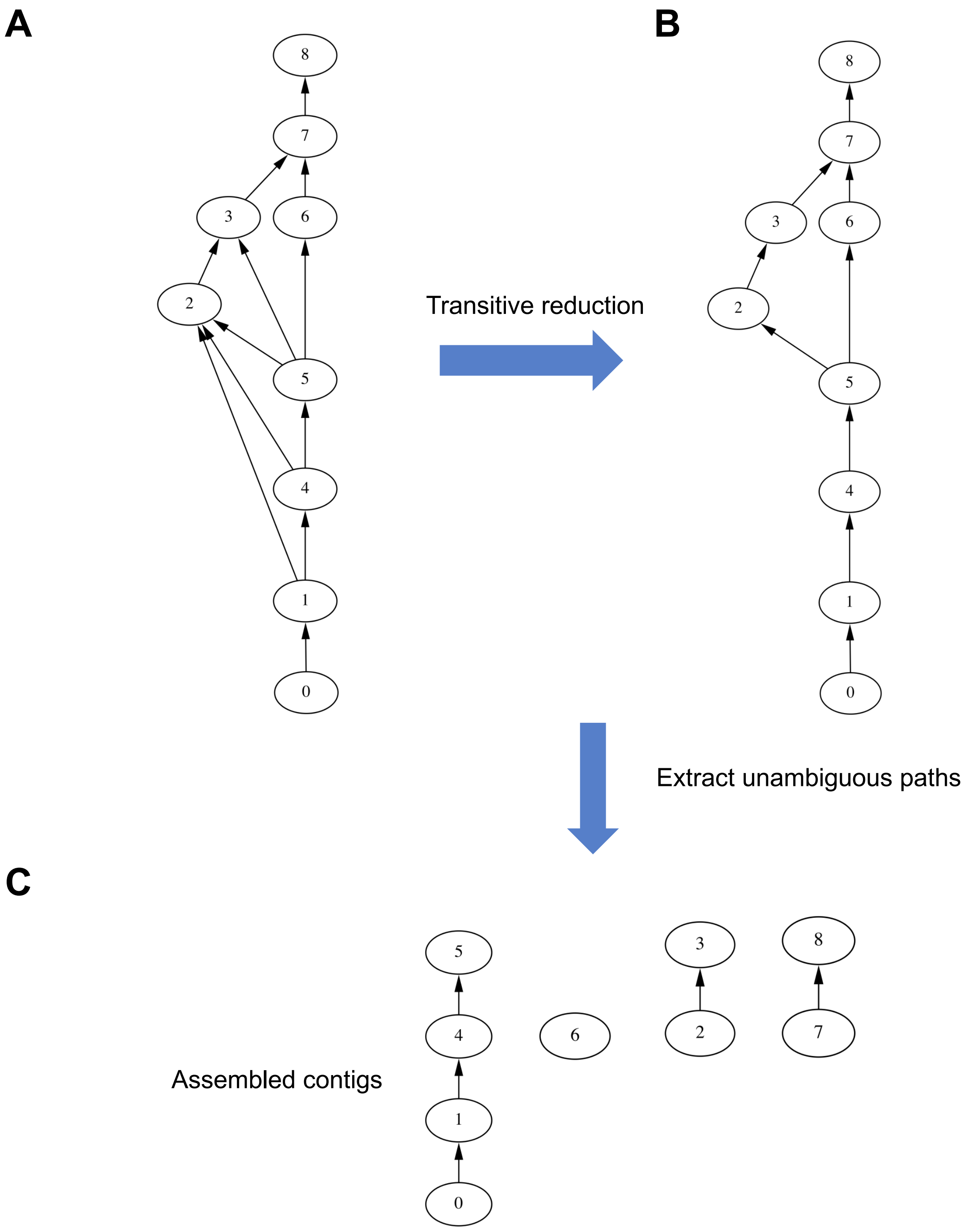


Figure S13: **Assembling draft contigs.** **A**, A demo graph obtained by overlapping draft contigs. Each vertex represents a draft contig and a edge means the two connected vertices overlap. **B**, Removing redundant edges by transitive reduction. **C**, Assembled contigs are obtained by extracting unambiguous paths from the graph in subfigure **B**.
